# SiRNA Molecules as Potential RNAi Therapeutics to Silence RdRP Region and N-Gene of SARS-CoV-2: An *In Silico* Approach

**DOI:** 10.1101/2022.10.08.511397

**Authors:** Mahedi Hasan, Atiya Tahira Tasnim, Arafat Islam Ashik, Md Belal Chowdhury, Zakia Sultana Nishat, Khandaker Atkia Fariha, Tanvir Hossain, Shamim Ahmed

## Abstract

COVID-19 pandemic keeps pressing onward and effective treatment option against it is still far-off. Since the onslaught in 2020, 13 different variants of SARS-CoV-2 have been surfaced including 05 different variants of concern. Success in faster pandemic handling in the future largely depends on reinforcing therapeutics along with vaccines. As a part of RNAi therapeutics, here we developed a computational approach for predicting siRNAs, which are presumed to be intrinsically active against two crucial mRNAs of SARS-CoV-2, the RNA-dependent RNA polymerase (RdRp), and the nucleocapsid phosphoprotein gene (N gene). Sequence conservancy among the alpha, beta, gamma, and delta variants of SARS-CoV-2 was integrated in the analyses that warrants the potential of these siRNAs against multiple variants. We preliminary found 13 RdRP-targeting and 7 N gene-targeting siRNAs using the siDirect V.2.0. These siRNAs were subsequently filtered through different parameters at optimum condition including macromolecular docking studies. As a result, we selected 4 siRNAs against the RdRP and 3 siRNAs against the N-gene as RNAi candidates. Development of these potential siRNA therapeutics can significantly synergize COVID-19 mitigation by lessening the efforts, furthermore, can lay a rudimentary base for the in silico design of RNAi therapeutics for future emergencies.

## Introduction

SARS-CoV-2, the cause behind the ongoing pandemic COVID-19, is a coronavirus strain under the virus family *Coronaviridae*. The positive sense single-stranded RNA genome (+ssRNA) of SARS-CoV-2 is almost 30 kb long, and the capsid contains four major types of structural proteins^1^. Just like all other coronaviruses, the 5’-terminus of the SARS-CoV-2 has the ORF1ab gene that encodes for non-structural proteins (nsp1 to nsp16), which are essential for viral replication, viral assembly, viral transcription and immune response modulation^2^. The 3’-terminus of the SARS-CoV-2 has open-reading frames (ORFs) that encode the four structural capsid proteins-spike proteins (S), envelope proteins (E), membrane proteins (M), nucleocapsid proteins (N), and also ORF3a, ORF6, ORF7a, ORF7b, ORF8, ORF10 genes^3^.

The SARS-CoV-2 ORF1ab encoded nsp12 is the RNA-dependent RNA polymerase (RdRP) that have vital functions in SARS-CoV-2 pathogenesis^2,4^. In coronaviruses, the RdRP is an enzyme that treats (+) ssRNA genome of RNA viruses as a template to produce the (-) RNA strand. This (-) RNA strand then guides viral transcription to produce sub-genomic mRNAs. Beside the transcription process, RdRP is also required for viral genome replication^5^. Hence, RdRP plays an indispensable role in the SARS-CoV-2 life cycle and can be an attractive target for novel antiviral therapeutics. Additionally, among the structural proteins of SARS-CoV-2, the N protein mediates the formation of helical ribonucleoproteins during the packaging of the RNA genome, thus plays a crucial role in viral assembly. It has a critical role in viral RNA replication and transcription by regulating viral RNA synthesis^6^. The N protein can modulate host cell metabolism and can suppress host innate immunity, too^7^. Therefore, the N protein can also be a suitable target for potential antiviral therapeutics.

A viral activity can be controlled by any means of silencing non-structural or structural genes. A gene can be silenced at different stages of infection, such as transcriptional, post-transcriptional or translational level^8–11^. Among the various mechanisms of gene silencing, RNA interference (RNAi) is one of them. The RNAi is a post-transcriptional gene silencing mechanism that either degrades or inactivates the messenger RNA (mRNA) of a corresponding gene with the help of some interfering RNA molecules, which are more or less 20-30 bp in their entire length^12–14^. Small interfering RNAs (siRNAs) are comparatively short noncoding double-stranded RNAs (dsRNAs) that are generally exogenous and designed experimentally. Although siRNAs are not encoded in the genome, but can be introduced in cellular cytoplasm with the help of nanoparticle vectors and various insertion techniques, which results in the degradation of homologous mRNA of the target gene^15^. It is now well-documented that siRNA can be efficient therapeutics because of its high target-specificity by perfect sequence complementarity with its target mRNA region and compatible insertion techniques into the target cell or organ^16^.

To summarize, *in vivo* siRNA insertion can either be achieved by classic viral delivery method or modern non-viral delivery methods^17–19^. Delivery of siRNA using viral vector is mediated by several retrovirus vectors, such as mouse stem cell virus (MSCV), the Moloney murine leukemia virus (MoMLV), and lentivirus. Also, adenovirus vectors are used to deliver siRNA into mammalian system^18,19^. on the other hand, the non-viral siRNA delivery system mainly comprised with five methods, for instance direct injection, cholesterol conjugation, lipid-based transfection, polymers and peptides delivery and delivery with surface modified nanoparticles^17^. High pressure induces siRNA penetration during direct injection^20^, cholesterol and lipid-based chemical agent (i.e., lipoplexes, polyplexes, and lipopolyplexes) mediated transfection are facilitated by lipoproteins used in those techniques^21,22^. The specificity and selectivity of delivery methods have increased with polymers and peptides mediated delivery as it can escape endosomal degradation^23^. Besides, surface-modified nanoparticle mediated siRNA delivery provides both protection and specificity^24^.

The introduced siRNA is then cleaved by RNase III type Dicer protein and becomes short double-stranded siRNA. The short antisense guide strand of the double-stranded siRNA is then recognized by cellular RNA-induced silencing complex (RISC). Whereas the sense passenger strand of siRNA is degraded by cellular enzymatic activity. The siRNA guide strand leads RISC to bind perfectly to the complementary mRNA, and the binding causes the cleaving and degradation of the mRNA by the Argonaute protein of RISC^25^. Recently, investigation of siRNAs is leading to the way of many antiviral defenses against several diseases, such as human immunodeficiency virus-1 (HIV-1)^26^, Dengue^27^, Influenza^28^, hepatitis C (HCV)^29^, and Zika^30^. The siRNAs are also being anticipated to treat various genetic disorders. As a result, the first global approval of a siRNA drug, named Patisiran, was announced by the U.S. Food and Drug Administration (FDA) in 2018^31^, to treat polyneuropathy in people with hereditary transthyretin-mediated amyloidosis. It is delivered into hepatocytes via lipid nanoparticles where it binds the 3′-UTR of both mutant and wild-type transthyretin (TTR) mRNA and causes its degradation, hence reducing TTR protein levels in serum and tissues^32^. Most recently in 2020, another siRNA drug has been approved by FDA, called Givosiran^33^, which is used to treat acute hepatic porphyria (AHP). The specific delivery of Givosiran is mediated by covalent linking of siRNA to a ligand and it targets aminolevulinate synthase 1 (ALAS1) mRNA, thus reduces the accumulation of δ-aminolevulinic acid and porphobilinogen successfully^34^.

In this study, the RdRP (nsp12) region and N gene of the SARS-CoV-2 viral genome were scanned to construct potential siRNA to silence these genes by utilizing some computational tools **(Figure 1)**. Thus, the predicted siRNAs, if developed will be capable of contributing to the fight against SARS-CoV-2 by degrading mRNA regions of RdRP and N. RNAi is going to be the next generation therapeutics because of its available insertion techniques, specificity, and cost-effectiveness^35^. Therefore, this study provides a benchmark for the establishment of RNAi therapeutics against SARS-CoV-2 as an effective treatment for COVID-19 as well as for any upcoming future pandemic situation.

**Figure 1:**
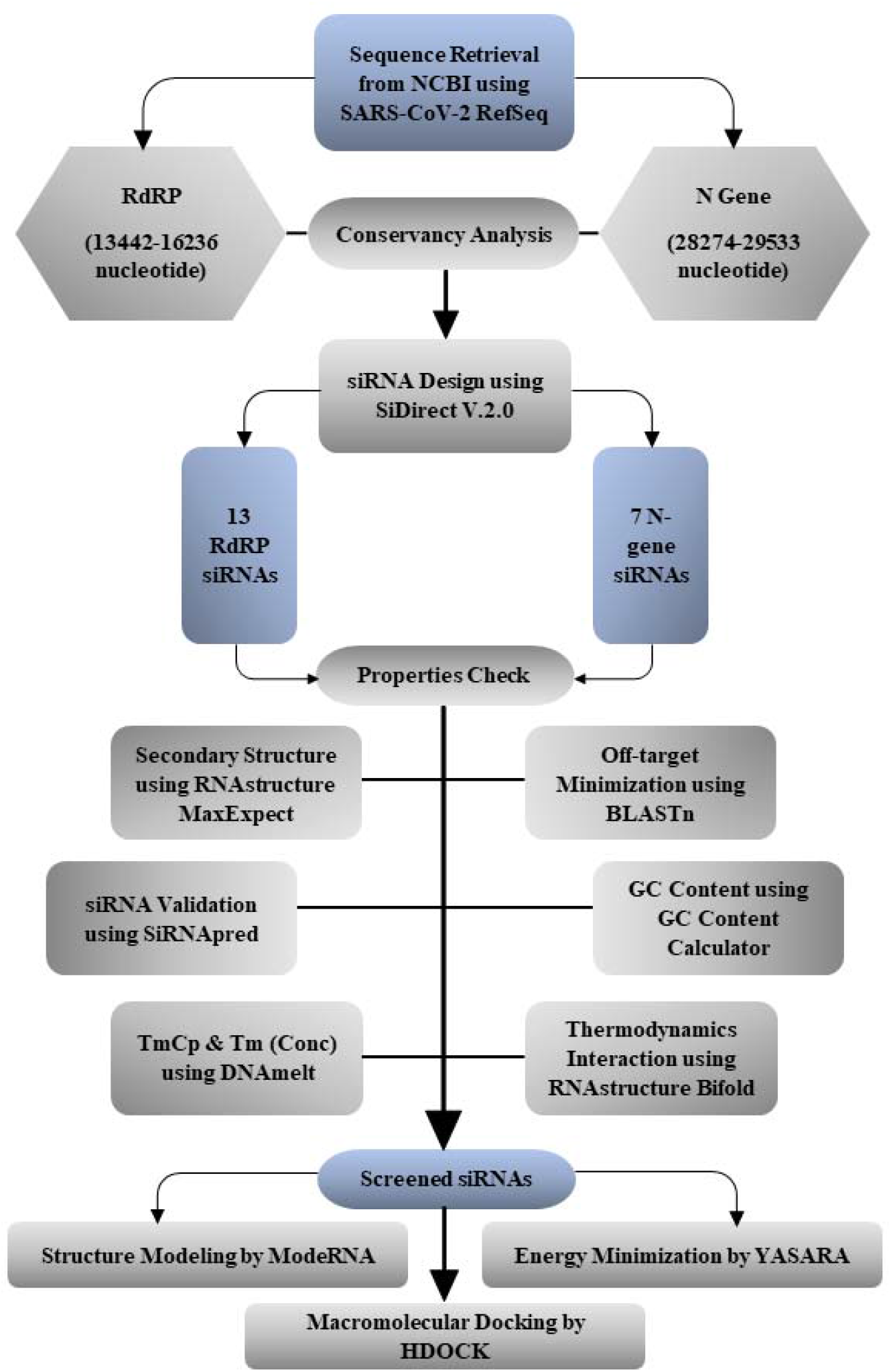
Graphical representation. Schematic diagram of siRNA prediction and optimization from the gene sequence.

## Result

### Conserved sequence among different variants

There were more than 97% similarities for RdRP and N among all available strains of the SARS-CoV-2 around the world. Also, the MEGAX integrated MSA derived all conserved sequences having at least 21 nucleotides are given in the **Supplementary table S2 and S3** for the RdRP and N, respectively. The conserved sequence positions were used to find the best siRNAs that will be effective against all reported variants of SARS-CoV-2. **Figure 2** represents all selected siRNA targets having 100% sequence conservancy among variants via the sequence logos.

**Figure 2:**
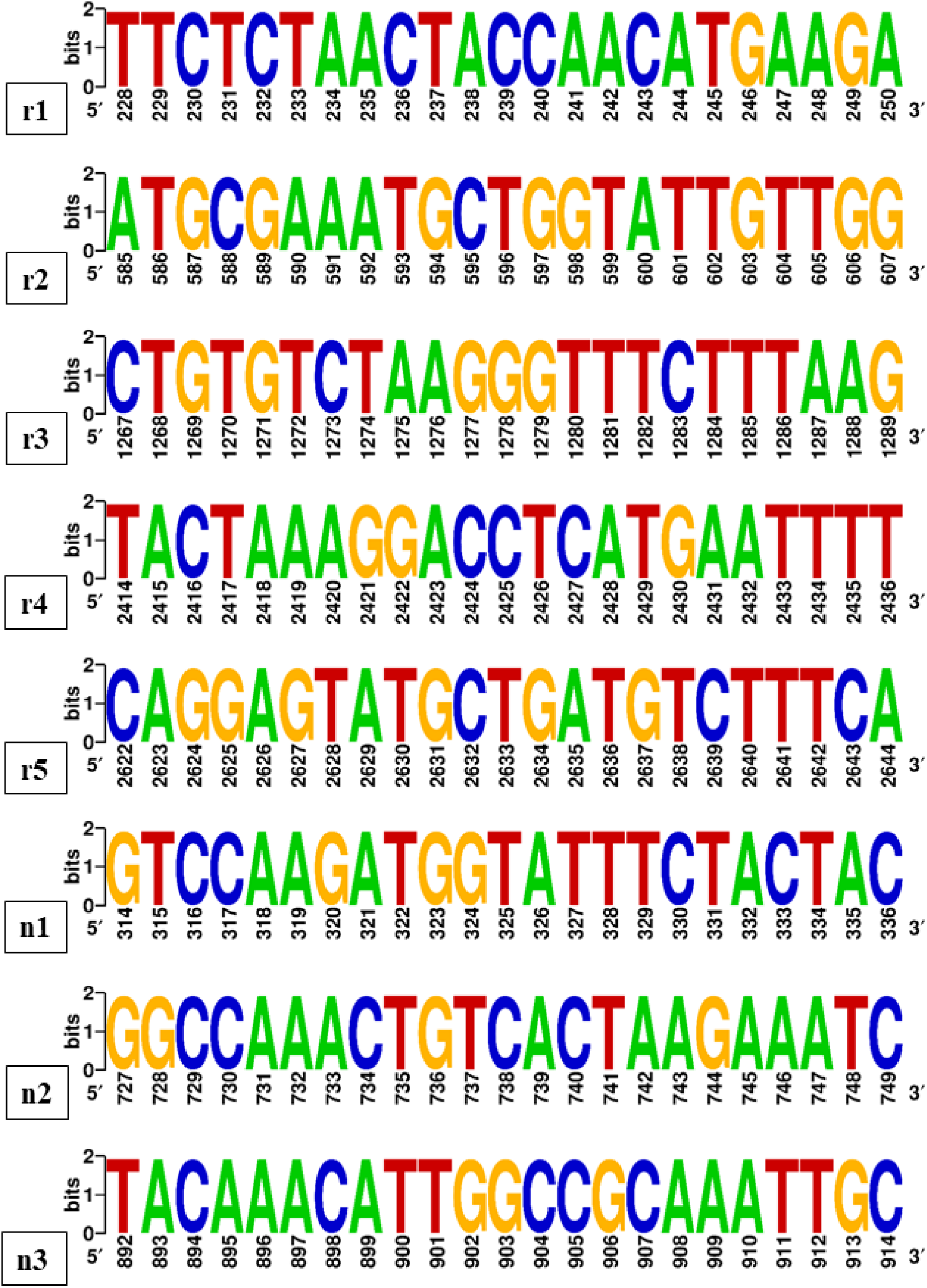
Conservancy analysis of selected siRNA targets for RdRP region and N gene. The RdRP and N sequences retrieved for MEGAX multiple sequence alignment are analyzed for conserved sequence of those two genes. The target sequences of screened siRNAs displayed 100% conservancy that has visualized using two bits long sequence logos constructed with WebLogo. The serial number below the logo indicates base position in RdRP and N separately. All bases of target sequences represent height of two bits long, hence producing 100% conservancy of corresponding bases in corresponding positions. rN, r = RdRP region targeting siRNAs, N = serial number of siRNAs; nN, n = N-gene targeting siRNAs, N = serial number of siRNAs.

### siRNA prediction with siDirect and minimized off-target

From the siDirect site, 13 siRNAs for RdRP region and 7 siRNAs for N-gene were found (**Supplementary tables S4 and S5**) to be potential on the basis of checked parameters. To find the T_m_ values for all predicted siRNAs of RdRP and N below 21.5 °C (**Supplementary tables S4 and S5**), the seed-target duplex was predicted, and all the siRNA duplexes were observed using the nearest-neighbor model and the thermodynamic parameters for the formation of siRNA duplexes. Also, there was no significant sequence similarity with the human genomic and transcript sequence database for the predicted siRNAs. Minimization of off-target sequence ensures more specifity for the siRNA target site complementation.

### Functional significance of GC content and RNA-RNA interaction by thermodynamics

The resultant GC content analysis using the GC content calculator showed an exact GC content percentage from 33.33% to 42.86% for the predicted siRNAs of both RdRP and N sequences (**Supplementary tables S4 and S5**), which is a suitable range for hybridization with a specific target^41,54^. Furthermore, the hybrid RNA structure of the predicted siRNAs and target sequences with their corresponding minimum free energy of hybridization (**Supplementary tables S4 and S5**) were measured using the RNA structure webserver’s bifold section, which is displayed in **Figure 3** for the selected siRNAs only.

**Figure 3:**
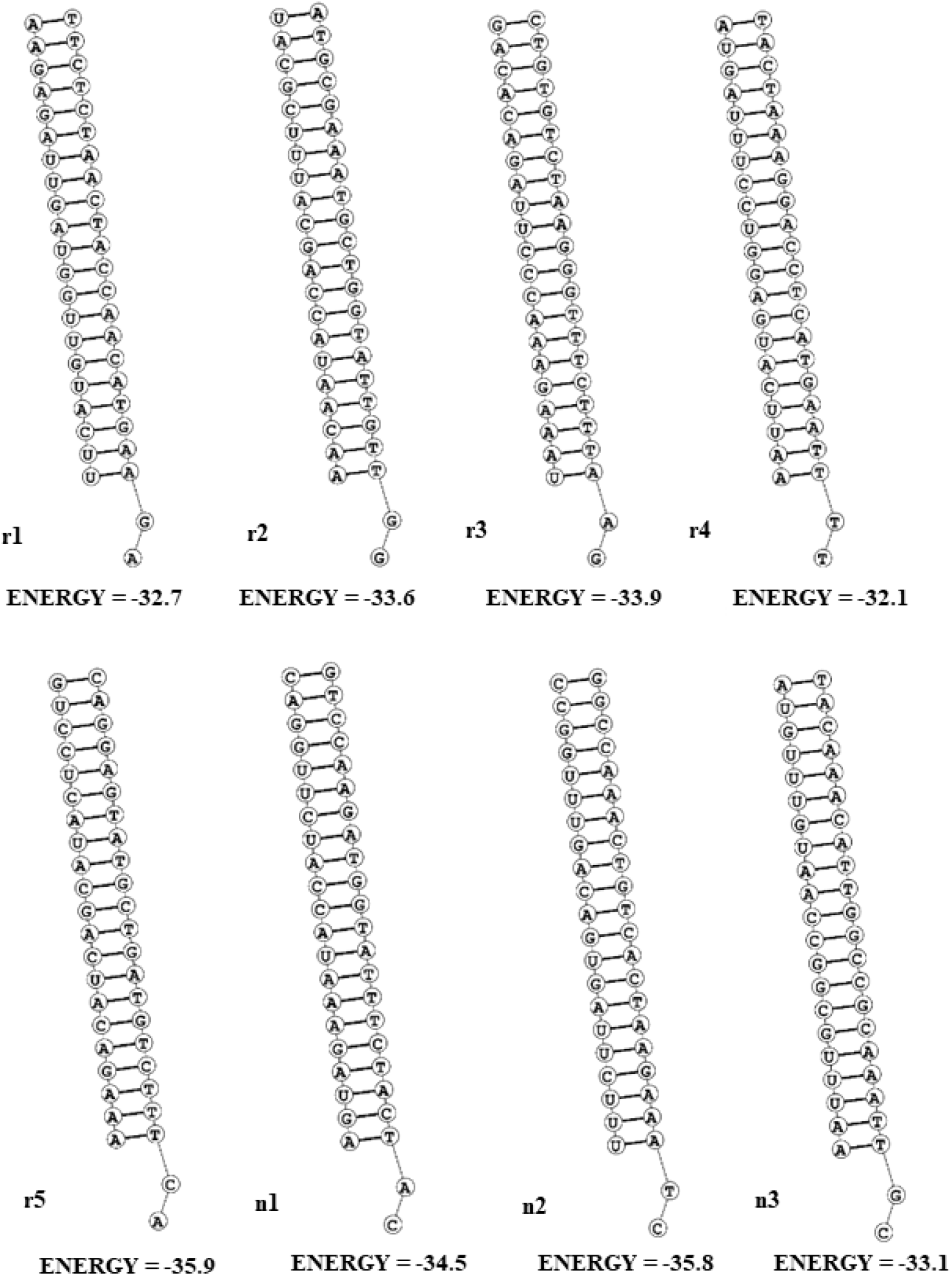
Secondary structures of potential target–siRNA duplex. The secondary structures of RdRP and N targeting siRNAs hybridized with their corresponding targets with their minimum free energy of hybridization (Kcal/mol). rN, r = RdRP region targeting siRNAs, N = serial number of siRNAs; nN, n = N-gene targeting siRNAs, N = serial number of siRNAs.

### Heat capacity and concentration plots of RNA-duplex

The higher values of T_m_C_p_ and T_m_(Conc) indicate higher effectiveness of siRNAs. These melting temperatures for predicted siRNAs of both RdRP and N were measured using the DNAmelt Server, where T_m_C_p_ and T_m_(Conc) values respectively for siRNAs of RdRP ranged from 79.9 °C to 85.1 °C and 79.0 °C to 83.7 °C (**Supplementary table S4**), and for siRNAs of N ranged from 80.7 °C to 85.5 °C and 80.1 °C to 84.4 °C (**Supplementary table S5**).

### Validation of predicted siRNAs

The validity and effectivity of the predicted siRNA molecules for both RdRP and N regions were measured by operating siRNAPred, and the siRNAs having validity (binary) values greater than 1 are considered as highly effective. Among 13 siRNA molecules for RdRP, 5 siRNAs were found to have value greater than 1, and among 7 siRNA molecules for N-gene, 1 siRNA had a value greater than 1. Besides, all other predicted siRNAs had validity (binary) values very close to 1 (**Supplementary tables S4 and S5**).

### Secondary structure prediction of siRNA guide strands

The positive values of minimum free energy of folding (value > 0) for siRNAs indicate better candidates because those siRNA molecules undergo least folding among themselves. To visualize the possible folding of the selected siRNA guide strands for both RdRP and N with their corresponding minimum free energy of folding (**Figure 4**) the RNA structure webserver was used. The values of minimum free energy of folding were in the range of 1.7–2.0 for RdRP siRNAs and of 1.4–1.8 for N siRNAs. The selected siRNAs showed unpaired terminal ends and less degree of folding for both RdRP region and N-gene.

**Figure 4:**
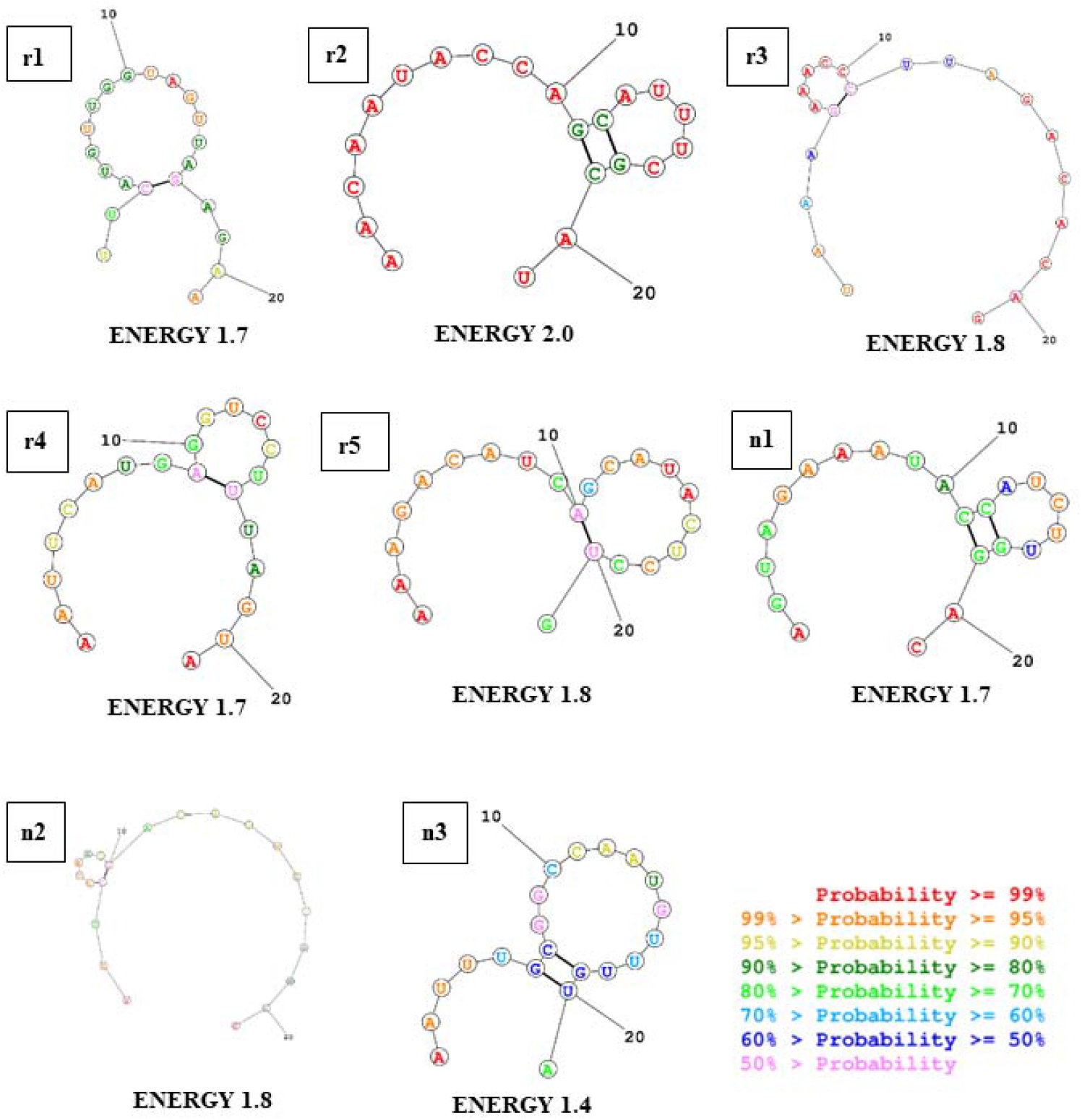
Secondary structures of siRNA guide strands. The values of minimum free energy of guide strands’ folding with visualization for both RdRP and N targeting siRNAs are illustrated and retrieved from RNA structure webserver. Color coding indicates the probability of predicted secondary structure positioning of that base. The probabilities with their corresponding colors are shown in inset. rN, r = RdRP region targeting siRNAs, N = serial number of siRNAs; nN, n = N-gene targeting siRNAs, N = serial number of siRNAs.

Finally, 5 siRNAs for RdRP and 3 siRNAs for N were selected as the best possible candidates against the SARS-CoV-2 based on different parameters, such as sequence conservation, GC percentages, T_m_ values, the free energy of hybridization, prediction validity, and free energy of folding of the siRNAs, and the T_m_ values for these selected siRNAs are given in **Table 1**.

**Table 1.**
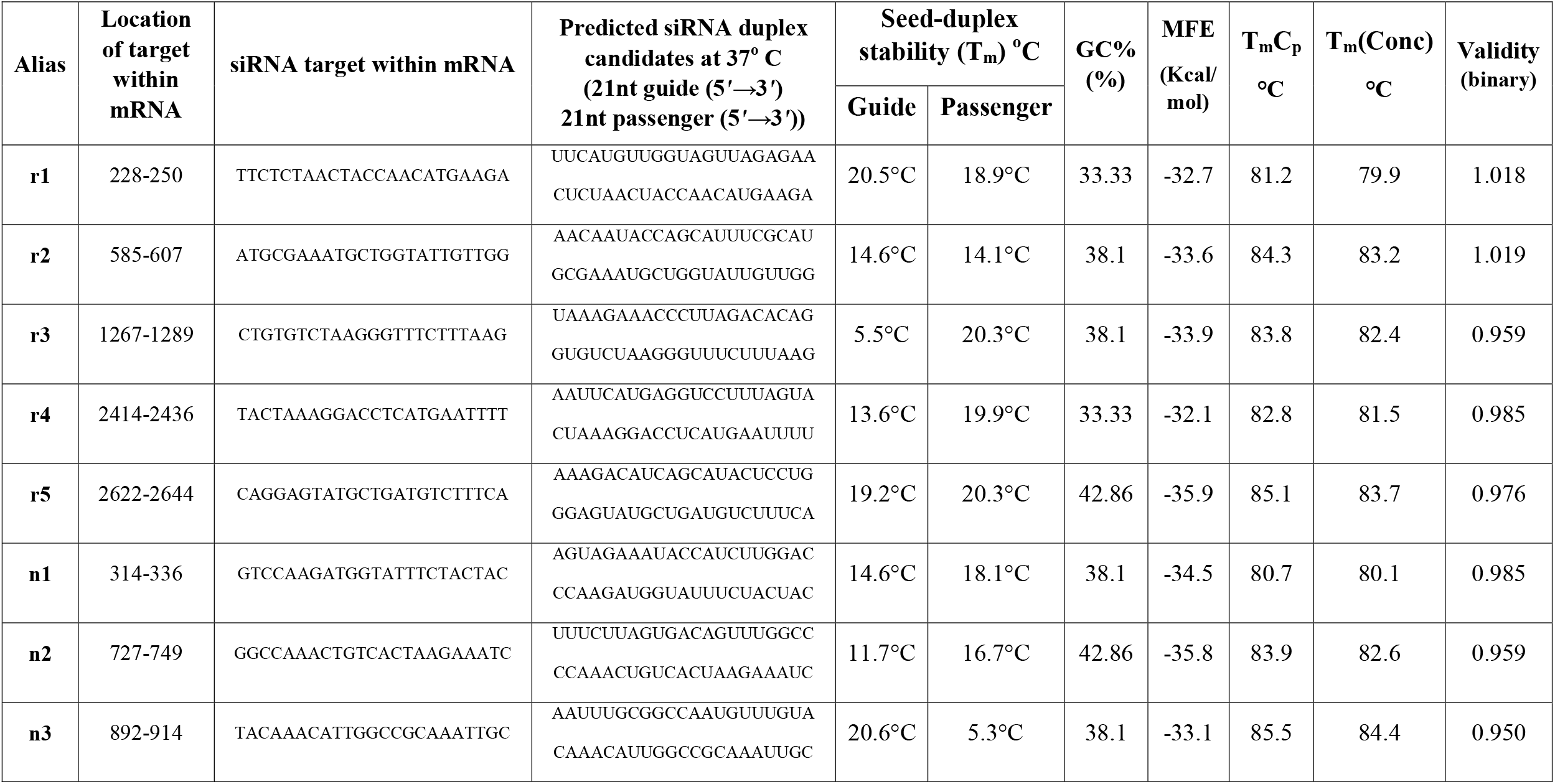
Effective siRNA molecules for RdRP and N regions with T_m_ values of both guide strand and passenger strand, guide strands’ GC%, free energy of binding (MEF) with target, T_m_C_p_, T_m_(Conc), and validity (binary). rN, N = serial number of RdRP targeting siRNAs; nN, N = serial number of N targeting siRNAs.

### 3D modeling and energy minimization of selected siRNAs

The 3D structures and energy-minimized structures of selected siRNA guide strands were modelled using ModeRNA and YASARA, respectively and visualized in PyMOL molecular graphics visualization software V.2.0 (**Figure 5**). The starting and ending energy values from YASARA energy minimization are reported in the **Table 2**.

**Table 2.** YASARA energy minimization values for screened siRNAs and the human Ago2 protein PAZ and MID domains interacting residues after docking with energy minimized siRNA molecules including their docking scores. rN, N = serial number of RdRP targeting siRNAs; nN, N = serial number of N targeting siRNAs.

| siRNAs | Energy minimization values |  | Docking score and interacting residues |  |  |
| --- | --- | --- | --- | --- | --- |
|  | Starting energy (kJ/mol) | Ending energy (kJ/mol) | Docking Score (Kcal/mol) | PAZ Domain | MID Domain |
| **r1** | 2004174.3 | -67729.5 | -313.37 | LYS266, ARG277, LYS278, TYR279, ARG280 | LEU522, GLY524, LYS525, THR526, TYR529, LYS533, THR544, GLN545, CYS546, VAL547, GLN548, ASN551, GLN558, THR559, ASN562, LEU563, LYS566, LYS570 |
| **r2** | 265228.7 | -57795.9 | -319.82 |  | LEU522, GLY524, LYS525, THR526, TYR529, GLN545, CYS546, VAL547, GLN548, GLN558, THR559, ASN562, LEU563, LYS566 |
| **r3** | -44569.7 | -75266.0 | -609.29 | ARG277, TYR279, ARG315, LYS335 | LEU522, GLY524, LYS525, THR526, TYR529, ALA530, LYS533, THR544, GLN545, CYS546, VAL547, GLN548, ASN551, GLN558, THR559, ASN562, LEU563, LYS566, LYS570 |
| **r4** | 166805899.4 | -63556.9 | -320.91 |  | GLN548, ASN551, ARG554, THR556, GLN558, THR559, ASN562, LEU563, LYS566 |
| **r5** | -32458.0 | -66374.0 | -315.42 | ALA227, LYS266, GLU268, LYS278, ARG280, VAL347 |  |
| **n1** | -35609.6 | -70222.5 | -625.91 |  | LEU522, PRO523, GLY524, LYS525, TYR529, LYS533, CYS546, VAL547, GLN548, LYS550 |
| **n2** | 286466.1 | -62300.3 | -412.41 | ARG280, GLN332, LYS335 | LEU522, GLY524, LYS525, TYR529, LYS533, GLN545, CYS546, VAL547, GLN548, LYS550, ASN551, GLN558, THR559, ASN562, LEU563, LYS566, LYS570 |
| n3 | 756245.5 | -67784.7 | -336.96 | ARG277, TYR279,<br>ARG315, GLN332 | LEU522, GLY524, LYS525, THR526, TYR529, LYS533,<br>GLN545, CYS546, VAL547, GLN548, ASN551, GLN558,<br>THR559, ASN562, LEU563, LYS566, LYS570 |

**Figure 5:**
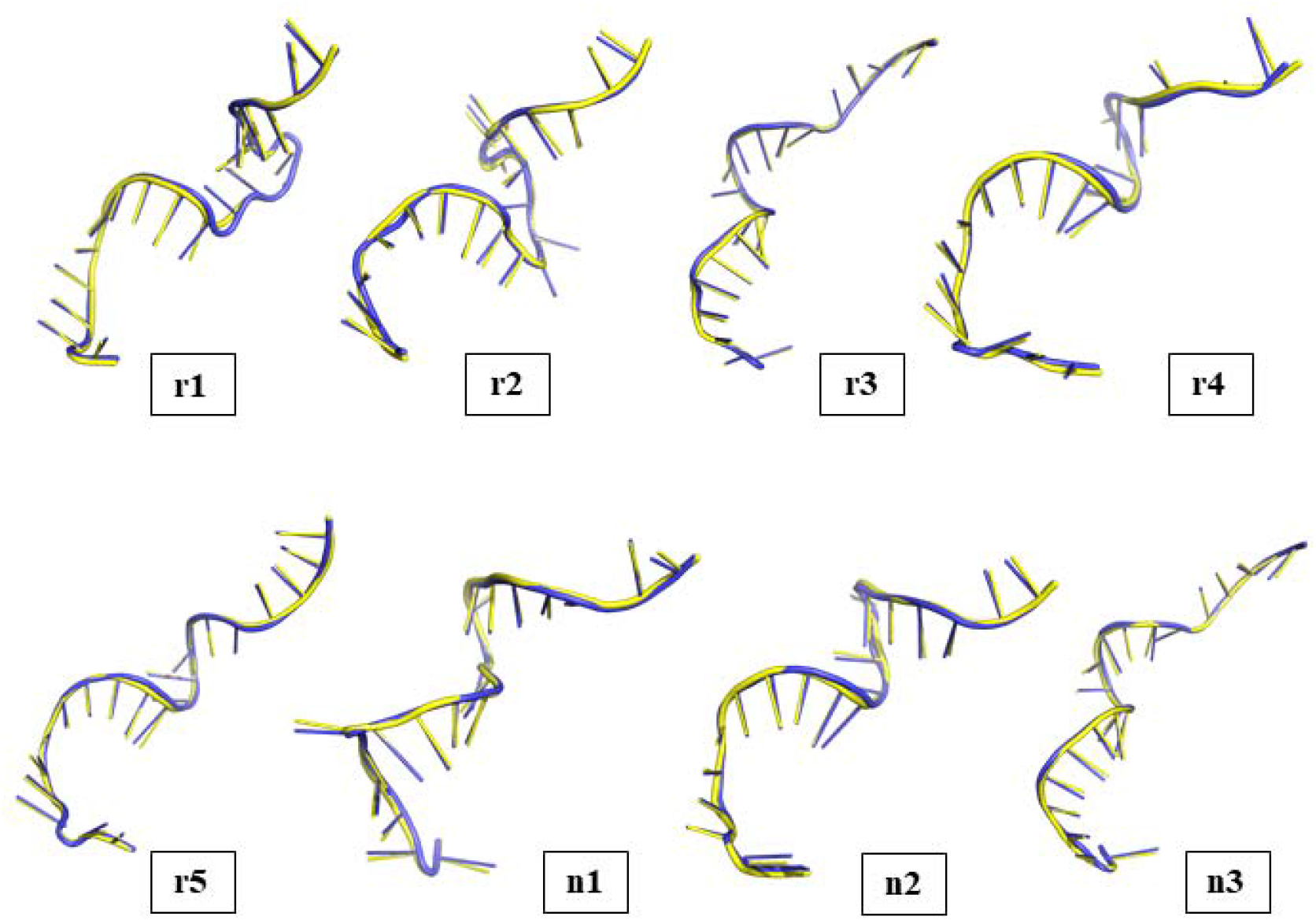
Three-dimensional (3D) structure visualization of YASARA energy minimized siRNAs. 3D secondary structure of siRNAs at starting energy (yellow color) in comparison with their energy-minimized versions (blue color), displayed using PyMol for both RdRP and N targeting siRNAs. rN, r = RdRP region targeting siRNAs, N = serial number of siRNAs; nN, n = N-gene targeting siRNAs, N = serial number of siRNAs.

### Macromolecular docking analyses

The HDOCK docking scores were in between -625.91 Kcal/mol to -313.37 Kcal/mol for the siRNAs n1 and r1, respectively. The docking model 1 given by the HDOCK server was the best docking model for all siRNAs, except the r2 siRNA, for which the best model was the HDOCK docking model 2. All of these docking performances showed effective Ago2-mediated mRNA silencing possibilities according to the Ago2-siRNA interaction pattern. Among them, the interaction patterns between Ago2 and r1, r3, n1, n2, n3 siRNAs were better than the remaining according to the interaction pattern. The HDOCK docking scores along with the PAZ and the MID domain interacting residues are given in the **Table 2**. The interacting residue-pair list is given in the **Supplementary table S6**. The docking results were finally visualized by PyMOL molecular graphics visualization software V.2.0 (**Figure 6**).

**Figure 6.**
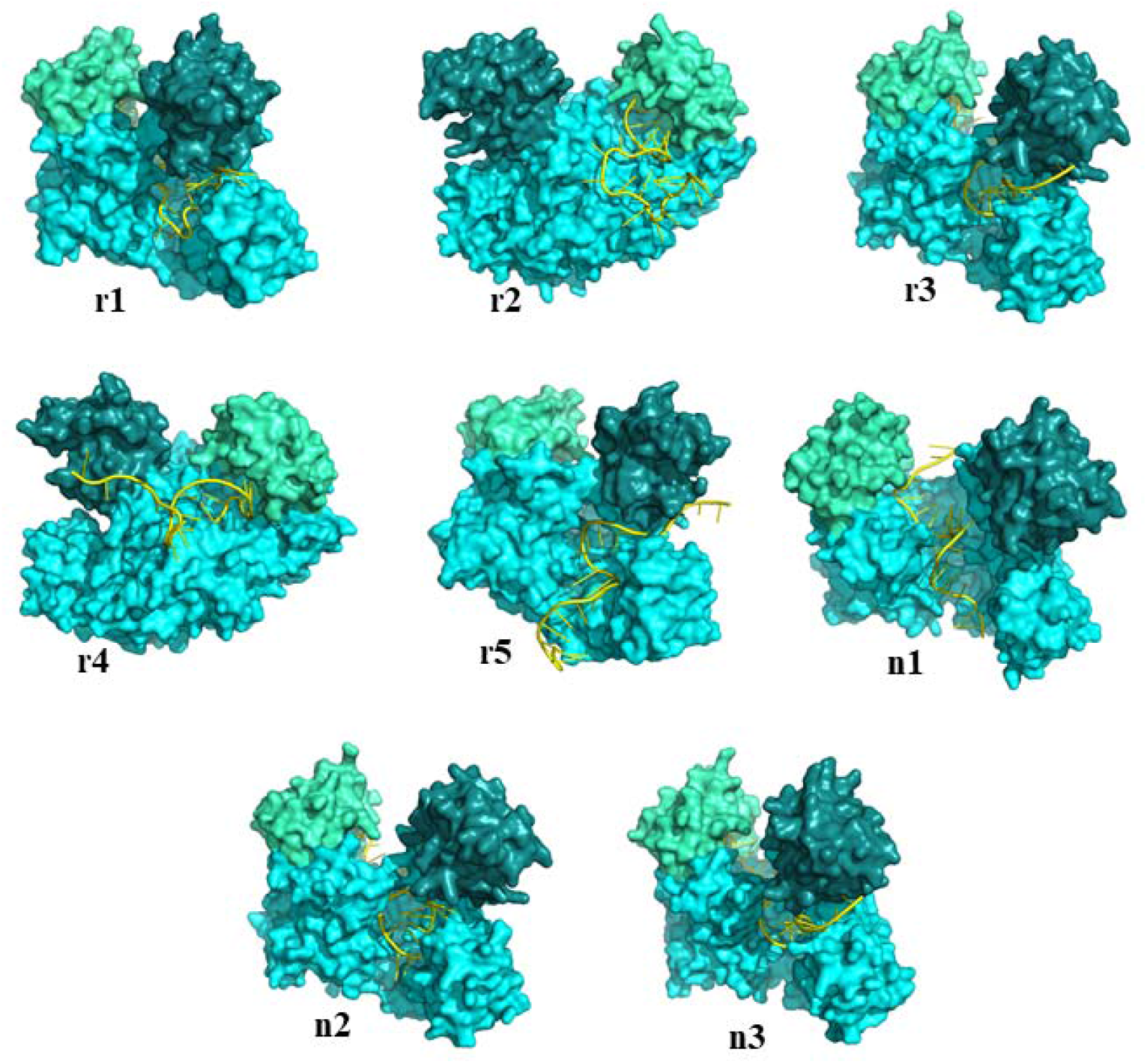
Visualization of argonaute protein and screened siRNAs binding interaction. Docking illustrations of siRNA (yellow cartoon view) with human Ago2 protein (cyan surface view), where the PAZ domain and the MID domain are colored as deep teal and greenish cyan, respectively. There siRNA-Ago2 docking interaction is visualized for potential siRNAs targeting the RdRP and N sequences of SARS-CoV-2. rN, r = RdRP region targeting siRNAs, N = serial number of siRNAs; nN, n = N-gene targeting siRNAs, N = serial number of siRNAs.

## Discussion

The ongoing global crisis, COVID-19, caused by the infection of SARS-CoV-2. According to World Health Organization (WHO) COVID-19 dashboard (https://covid19.who.int/), more than 500 million human respiratory illnesses, including more than six million deaths caused by this disease till this year. Sustainable and effective treatment regimen against COVID-19 along with proper confinement strategy against its causal agent is still beyond the reach, calling the need for potential therapeutics to fight the pandemic. The present study was carried out using the RdRP and N sequences of the SARS-CoV-2 to design siRNA molecules as potential antivirals against COVID-19. The RdRP is a non-structural protein (nsp12) encoded in the ORF1ab gene, which is required for the replication, transcription, and increasing the copy number of the SARS-CoV-2 genome^55^. Whereas the N-gene encodes nucleocapsid phosphoprotein, which is required for viral RNA genome packaging, assembly, and interaction with host metabolism or immunity^56^. Thus, the RdRP and N sequences are suitable targets for silencing via siRNAs, thereby disrupting SARS-CoV-2 life cycle.

The BLAST result between all available SARS-CoV-2 strains in NCBI database showed more than 97% similarities for RdRP and N sequences. Also, the conservancy analysis between different variants of SARS-CoV-2, such as the alpha variant, beta variant, gamma variant and delta variant showed highly conserved sequence similarities for these aforementioned genes. The RdRP region and N-gene were 99.21% and 97.14% conserved respectively (**Supplementary tables S2 and S3**) for all stated variants that was confirmed by MEGAX multiple sequence alignment program. Later, the conservancy of screened siRNA targets were visualized in **Figure 2** as two bits long sequence logos. A sequence logo is a 2D graph having base position in x-axis and bits in Y-axis. Base frequency is the measurement of how frequent a specific nucleotide is present in each position of an MSA. This base frequency is used to measure the entropy of that position and subsequently the height (Bits - Entropy) of all bases of that corresponding position. The most frequent base in a particular position has least entropy, hence highest height. Thus, bases having height equals to bits are 100% conserved^57^. According to **Figure 2**, all base position of every siRNA target sequences had complete 2 bits long base heights, which indicates 100% sequence conservancy among all variants of SARS-CoV-2 in these particular positions.

The siRNA molecules are short noncoding exogenous RNA molecules that can effectively silence any specific gene by degrading its corresponding mRNA via perfect complementarity between the siRNA guide strand and the target mRNA. The siDirect 2.0 software predicted possible siRNAs for the target sequences and distinguished off-target-free seed sequences properly from the off-target-positive ones^40^ because the T_m_ of 21.5 °C was marked while running the program. A total of 13 siRNAs for the RdRP sequence and 7 siRNAs for the N-gene sequence were found from the siDirect site for further analysis. Although the siDirect 2.0 software can minimize the off-target for designing siRNAs, the NCBI BLASTn similarity search was executed for all predicted siRNAs, and no significant sequence similarities were found with the human genomic plus transcript database.

The siRNAs with high GC% can interfere with siRNA duplex unwinding and effective degradation of mRNA targets. Also, too low GC content can decrease target mRNA recognition and hybridization with the target^41^. Hence, the suitable GC range is an essential parameter for siRNA’s proper functioning^54^. In this present study, an exact GC content percentage from 33.33% to 42.86% was found using the GC content calculator for the predicted siRNAs of both RdRP and N, which is within a suitable range to support the hybridization of those siRNAs with their specific target mRNAs^41,54^. This range was also maintained by similar previous studies, for instance the study of *Chowdhury U*.*F. et al (2021)*^58^ and *Shohan M*.*U*.*S. et al (2018)*^59^. Also, the hybridization and minimal free energy of binding (Kcal/mol) with target for the predicted siRNAs were -32.7, -33.6, -33.9, -32.9, -31.7, -32.5, -35.2, -33.5, -34.5, -32.1, -35.9, -32.7, -34.3 for RdRP and -34.5, -34.2, -35.8, -34.5, -32.5, -33.1, -31.7 for N, which were found using the bifold section of RNA structure webserver (**Supplementary tables S4 and S5**) as well as their corresponding hybridization secondary structures for selected siRNAs are illustrated in **Figure 3**. The more negative free energy of binding value, the more strong binding of the predicted siRNAs with their target sites^60^, and therefore the predicted siRNAs can also suitably bind to their targets.

Moreover, heat capacity plot and concentration plot are crucial benchmarks to predict the siRNA thermodynamics interactions. In the heat capacity plot, the heat capacity (C_p_) is plotted as a function of temperature, and when it is a function of melting temperature (T_m_) it is called T_m_(C_p_), which is the local maximum of the heat capacity curve. On the other hand, in the concentration plot, molar fractions are plotted as a function of temperature, and concentration as a function of T_m_ is T_m_(Conc), which is the point where the concentration of the siRNA duplex is one-half of its maximum value^47^. The DNAMelt web server was run to find the T_m_C_p_ and T_m_(Conc) values, and the higher values lead to the best siRNAs as found in some similar study, for example the study of *Nur S*.*M. et al (2015)*^61^ and *Shohan M*.*U*.*S. et al (2018)*^59^. In this study, the predicted siRNAs showed high melting profiles, as shown in **Table 1**, hence regarded as the best candidates as potential antivirals.

Additionally, primarily selected siRNA molecules were validated to measure their efficacy, and siRNAs were selected as potential antiviral candidates for their high validity score (value ∼1 or ≥ 1) (**Table 1**)^58,62^. As crystal structures of RNAs are not available, using computational methods to design RNA secondary structures is much valuable^63^. The MaxExpect program determined the predicted siRNAs’ secondary molecular structures by computational methods in the RNA structure web server, and the secondary folding structure of selected siRNAs are shown in **Figure 4**. An unstructured guide strand of siRNA can easily bind with the target mRNA and silence it. Nevertheless, a base-paired terminal of the siRNA guide strand is less available to bind with the complementary mRNA and becomes inactivated^64^. Hence, if the degree of guide RNA folding structure decreases, the gene silencing by siRNA becomes much more efficient. In these analyses, the selected siRNAs also displayed the availability of nucleotides in the terminal regions and less folding degree (**Figure 4**). Finally, 5 siRNAs for RdRP region and 3 siRNAs for the N-gene were selected as suitable antiviral candidates for effective post-transcriptional gene silencing of SARS-CoV-2 based on different measured parameters.

In the RNAi pathway, siRNA molecules are intrinsically double-stranded that eventually separate, and only the guide strand interacts with RISC^65,66^. As the guide strand plays a prominent role in this pathway, it was inevitable to focus on the guide strand’s energy minimization. The energy minimization is one of the crucial steps to measure the compatibility of a guide strand during *in vivo* functions^67–69^. If the potential energies of the guide strand’s 3D models would much higher, they would not be compatible with the living system. The energy minimization process can resolve improper structures and stereochemistry, can add ions and water molecules, thus subsequently can relax the system and re-position atoms’ arrangement in molecules at its local minima using the principle of Molecular Mechanics (MM)^70,71^. In this study, the YASARA force field was used for all residues, and RNA atom type description was included in the parameter file. The lower ending potential of each strand indicates that the structure has minimized (**Table 2**), and favors the *in vivo* functioning^72^.

Macromolecular docking between human Ago2 protein and energy minimized modelled siRNAs shows perfect interaction pattern, specially, with the PAZ and MID domain of Ago2 (**Table 2**). Generally, both ends of an siRNA should be anchored to exert an effective silencing, the PAZ domain interacts in a weaker binding manner with the 3′-end of siRNAs, and the MID domain strongly interacts with the 5′-end of the siRNAs. Because, the 3′-end of an siRNA needs to be switched between a bound and unbound manner during the nuclease activity of Ago2 protein^73–75^. In this study, among the selected 8 siRNAs, 4 siRNAs namely r1, r3, n2, n3 have shown effective binding with both PAZ and MID domain. Whereas another 3 siRNAs namely r2, r4, and n1 have shown interaction only with the MID domain. But an exceptional case is observed for r5, which only interacts with the PAZ domain of Ago2.

According to the surface residue interaction patterns from this macromolecular docking (given in **Supplementary table S6** and visualized in **Figure 6**) the siRNAs, namely r1, r3, n1, n2 and n3 might be the best siRNAs since they have shared interacting residues with some wet lab interaction studies^73,76,77^. All these 5 siRNAs contain stably interacting arginine (ARG) residues, such as the ARG179, ARG635, ARG710, ARG761, ARG792, and 4 of them have ARG812 interacting residue (except n1) too. Although these residues are PAZ and MID domain interface residues, they are verified by the studies mentioned above. Except the r1 siRNA, the other stated 4 siRNAs have TYR804 interaction with Ago2, which was also found in the aforementioned studies. Another 2 siRNAs r2 and r4 also have shown ARG792 interaction but the interaction pattern for r5 is, however, poor in manner and can be negligible.

Among the PAZ domain interacting residues, ARG277, TYR279 are the most common and for the MID domain interacting residues GLN548 interaction is the most common, which is observed in r1, r2, r3, r4, n1, n2, n3. Some other MID domain interactions include GLN548, GLN558, THR559, ASN562, LEU563, LYS566 that have been observed for 6 aforementioned siRNAs except the n1 and the LEU522, GLY524, LYS525, TYR529, CYS546, VAL547 interactions are found in other 6 aforementioned siRNAs except the r4. All of these interaction are common and verified by other studies stated above^73,76,77^. As shown in the **Supplementary table S6**, the r5 siRNA has an unusual interaction pattern and becomes insignificant. But the interaction patterns for the other 7 siRNAs cannot be neglected, hence it can be said that they hold promising potential to effectively silence their corresponding target RdRP and N mRNA sequences of SARS-CoV-2.

The HDOCK docking scores for siRNAs targeting RdRP region range from -609.29 Kcal/mol to -313.37 Kcal/mol that correspondence to r3 and r1, respectively (**Table 2**). Also, the HDOCK docking scores for siRNAs targeting N-gene range from -625.91 Kcal/mol to -336.96 Kcal/mol that respectively corresponding to n1 and n3 (**Table 2**). These scores are much favorable for all of these selected siRNAs and indicate their strong interaction with the human Ago2 protein. Although the interaction patterns are similar to previously mentioned verified studies^73,76,77^, it leaves further confirmation about the effectiveness of these siRNAs via further *in vitro* and *in vivo* studies.

From our previous study, both miRNAs and siRNAs were predicted against the SARS-CoV-2 ORF1ab gene^78^. Another recent study^58^ had shown an almost similar strategy targeting the N and S protein gene of SARS-CoV-2. As this current study targets the most crucial protein of SARS-CoV-2, the RdRP, to identify and design best siRNAs against it, as well as identified 3 potential N-gene siRNAs, this study can also add some valuable inputs in the race of applying therapeutics against SARS-CoV-2. Also, inclusion of most recent delta variant genome and other variants of SARS-CoV-2 makes these predicted siRNAs relatively more effective against SARS-CoV-2 in comparison with other previous studies.

Currently, evidence has suggested that *in silico* assessment might be a novel and feasible approach for viral therapeutics development. In this study, 07 siRNA molecules were predicted as the credible candidates against RNA-dependent RNA polymerase and nucleocapsid phosphoprotein gene of SARS-CoV-2 using all maximum parameters in optimum conditions including their binding interaction analysis with argonaute protein. Additionally, these systematic analyses will provide a guideline for the future establishment of a computational model to anticipate siRNA antiviral therapeutics. Moreover, the compatible siRNA insertion techniques (via lipid-based transfection, nanoparticle, etc.) into the target cells or organs^79^, RISC-mediated target specific mRNA degradation, and cost-effective manner^35^ warrants the siRNA therapeutics an effective treatment for the ongoing pandemic, COVID-19. Besides, the workflow of this study will guide the *in silico* design of RNAi therapeutics and also will inspire researchers for developing RNAi therapeutics to control any upcoming pandemic situation caused by infectious agents faster.

## Materials and methods

### Sequence retrieval and analysis

The reference sequence of SARS-CoV-2 from the National Center for Biotechnology Information (NCBI), GB: NC_045512.2, was used to obtain the target sequences “RdRP or nsp12” and “N”. The NCBI graphics database was used to retrieve the nucleotide sequence of 13442-16236 bp for RdRP and of 28274-29533 for N (https://www.ncbi.nlm.nih.gov/nuccore/NC_045512.2?report=graph). After that, the nucleotide sequences were utilized to perform the BLAST (Basic Local Alignment Search Tool) (https://blast.ncbi.nlm.nih.gov/Blast.cgi) to achieve their sequence similarities among all available strains of the SARS-CoV-2 around the world in GenBank, the National Institutes of Health (NIH) genetic sequence database^36^.

### Sequence conservancy analysis among variants

Nearly 50 recent completely sequenced SARS-CoV-2 genome from all 6 geographical continents around the globe for the early pangolin lineages, A and B, was derived from NCBI Virus SARS-CoV-2 resources (https://www.ncbi.nlm.nih.gov/labs/virus/vssi/#/sars-cov-2) and subsequently the RdRP and N sequences were retrieved from them. Meanwhile, each geographic continent that had at least one complete sequence of SARS-CoV-2 variants, such as the alpha variant, the beta variant, the gamma variant, and the delta variant sequences having pangolin lineages B.1.1.7, B.1.351, P.1, and B.1.617.2, respectively were retrieved by using the reported website for the RdRP and N sequences of these variants. Later, evolutionary analysis by maximum likelihood method^37^ were performed by using the MEGAX software^38^ integrated multiple sequence alignment (MSA) program for 60 different sequences. Finally, the conservancy was visualized as sequence logos for selected RdRP and N siRNA target sequences that was constructed by the WebLogo webserver^39^.

### siRNA prediction and off-target minimization

To predict potential siRNAs for target nucleotide sequences, RdRP and N, an online web server siDirect version 2.0^40^ was operated. Several parameters, including melting temperature below 21.5 °C, GC content 31.6%-57.9%^41,42^ and Ui-Tei^43,44^, Renold^45^, Amarzguioui^41^ combined rules (algorithms have given in **Supplementary table S1**) were checked in the siDirect site. For the formation of a stable RNA duplex the seed-target duplex stability (T_m_) was also calculated. Consequently, nucleotide BLAST (BLASTn) program was executed for siRNA target sequences using the human genomic plus transcript database by default parameters to check off-target sequences in the genome of any non-targeted sites for more specific siRNA selection.

### GC content calculation and measurement of thermodynamic interaction

GC Content Calculator (http://www.endmemo.com/bio/gc.php) was operated to calculate the exact GC content percentages in predicted siRNAs. Afterwards, the thermodynamics interaction between the predicted viral siRNAs and the target sequences was measured by the RNA structure web server’s bifold section (http://rna.urmc.rochester.edu/RNAstructureWeb/Servers/bifold/bifold.html).

### Calculation of the heat capacity and concentration plot

The melting temperature T_m_C_p_, heat capacity curve’s local maximum, and T_m_(Conc), one-half concentration of maximum value of a double-stranded molecule were measured by operating the hybridization of two different strands option (http://unafold.rna.albany.edu/?q=DINAMelt/Hybrid2) of the DNAmelt Server^46,47^ with default values and in simple form checking the RNA option for the predicted siRNAs.

### Validation of predicted siRNA molecules

The predicted siRNA molecules were screened using the Main21 dataset with the support vector machine algorithm and by the binary pattern prediction approach to measure the actual efficacy of the siRNA molecules and their validation via operating an online free accessible server siRNApred (http://crdd.osdd.net/raghava/sirnapred/).

### Secondary structure prediction

The MaxExpect^48^ program in the RNA structure web server^49^ was operated to anticipate the secondary structure of siRNA guide strands with their respective free energy of folding.

### Modeling the 3D structures of predicted siRNAs and Energy Minimization

For validating predicted siRNAs’ function *in vivo*, it is indispensable to construct the 3D structure of siRNAs and minimize energy. Construction of a three-coordinated arrangement of guide strand was done using template-based sequence alignment followed by exporting the 3D model in PDB file format via an advanced online tool for modeling RNA 3D structures, the ModeRNA server^50^. For this purpose, the guide RNA strand bound to the human Argonaute2 (Ago2) protein (PDB id: 4W5Q) was used as a template. In searching for the lowest potential conformation (local minima), the energy was further minimized using YASARA Energy Minimization Web Server^51^. YASARA Energy Minimization Web Server uses YASARA force field. The YASARA force field includes bonded parameters (bond length, bond angle, and dihedrals) and non-bonded parameters (electrostatic interactions and Lennard-Jones potentials) for corresponding atom types of RNA molecules^52^.

### Macromolecular docking of Ago2 and predicted siRNAs

The macromolecules for conducting an effective RNAi silencing, the human Ago2 protein and predicted siRNAs, were docked by using the HDOCK server (http://hdock.phys.hust.edu.cn/) to investigate their interaction pattern with each other. HDOCK is an online server that perform any macromolecular docking based on a hybrid algorithm of template-based modeling and ab-initio free docking system^53^. The receptor protein Ago2 was derived from RCSB Protein Data Bank (https://www.rcsb.org/) (PDB id: 4W5Q) and refined to extract the Ago2 protein from the PDB cluster. Whereas the ligands were YASARA energy-minimized siRNAs in PDB format. Based on the best docking scores and the protein-siRNA interaction pattern, the best docking performances were evaluated.

## Data availability statement

We have used publicly available free databases as the main data source that are mentioned in method section with proper referencing. The supplementary data file contains all produced data during analyses.

## Acknowledgements

The authors would like to acknowledge the Department of Biochemistry and Molecular Biology, Shahjalal University of Science and Technology, Sylhet for laboratory facilities and logistic supports.

## Author contributions

Study design: M.H., A.T.T. and A.I.A.; Conceptualization: S.A. and T.H.; Data curation: M.H.,

A.T.T. and A.I.A.; Experimentation: M.H., A.T.T., A.I.A. and Z.S.N.; Result interpretation: M.H., A.T.T., A.I.A. and Z.S.N.; Figure representation: A.T.T. and A.I.A.; Manuscript writing: M.H., A.T.T., M.B.C. and Z.S.N.; Critical reviewing: M.B.C., K.A.F., T.H. and S.A. All authors provided critical feedback and approved the final manuscript.

## Additional Information

The authors declare no competing interests among them.

## Notes

### Competing Interest Statement

The authors have declared no competing interest.

## References

1. Wu, F. et al. Complete genome characterisation of a novel coronavirus associated with severe human respiratory disease in Wuhan, China. bioRxiv 2020.01.24.919183 (2020) doi:10.1101/2020.01.24.919183.

2. Wu, F. et al. A new coronavirus associated with human respiratory disease in China. Nature 579, 265–269 (2020).

3. Prajapat#, M. et al. Drug targets for corona virus: A systematic review. Indian J. Pharmacol. 52, 56 (2020).

4. Ogando, N. S. et al. SARS-coronavirus-2 replication in Vero E6 cells: replication kinetics, rapid adaptation and cytopathology. J. Gen. Virol. 101, 925–940.

5. Wang, Y., Anirudhan, V., Du, R., Cui, Q. & Rong, L. RNA-dependent RNA polymerase of SARS-CoV-2 as a therapeutic target. J. Med. Virol. n/a,.

6. Dutta, N. K., Mazumdar, K. & Gordy, J. T. The Nucleocapsid Protein of SARS–CoV-2: a Target for Vaccine Development. J. Virol. 94, (2020).

7. Li, J.-Y. et al. The ORF6, ORF8 and nucleocapsid proteins of SARS-CoV-2 inhibit type I interferon signaling pathway. Virus Res. 286, 198074 (2020).

8. Curradi, M., Izzo, A., Badaracco, G. & Landsberger, N. Molecular Mechanisms of Gene Silencing Mediated by DNA Methylation. Mol. Cell. Biol. 22, 3157–3173 (2002).

9. Filipowicz, W., Jaskiewicz, L., Kolb, F. A. & Pillai, R. S. Post-transcriptional gene silencing by siRNAs and miRNAs. Curr. Opin. Struct. Biol. 15, 331–341 (2005).

10. Park, Y.-D. et al. Gene silencing mediated by promoter homology occurs at the level of transcription and results in meiotically heritable alterations in methylation and gene activity. Plant J. 9, 183–194 (1996).

11. Shaklai, S., Amariglio, N., Rechavi, G. & Simon, A. J. Gene silencing at the nuclear periphery. FEBS J. 274, 1383–1392 (2007).

12. Fire, A. et al. Potent and specific genetic interference by double-stranded RNA in Caenorhabditis elegans. Nature 391, 806–811 (1998).

13. Hannon, G. J. RNA interference. Nature 418, 244–251 (2002).

14. McManus, M. T. & Sharp, P. A. Gene silencing in mammals by small interfering RNAs. Nat. Rev. Genet. 3, 737–747 (2002).

15. Kanasty, R., Dorkin, J. R., Vegas, A. & Anderson, D. Delivery materials for siRNA therapeutics. Nat. Mater. 12, 967–977 (2013).

16. Chen, P. Y. et al. Strand-specific 5′-O-methylation of siRNA duplexes controls guide strand selection and targeting specificity. RNA 14, 263–274 (2008).

17. Gao, K. & Huang, L. Nonviral Methods for siRNA Delivery. Mol. Pharm. 6, 651–658 (2009).

18. Rozema, D. B. & Lewis, D. L. siRNA delivery technologies for mammalian systems. TARGETS 2, 253–260 (2003).

19. Sioud, M. On the delivery of small interfering RNAs into mammalian cells. Expert Opin. Drug Deliv. 2, 639–651 (2005).

20. Takei, Y., Kadomatsu, K., Yuzawa, Y., Matsuo, S. & Muramatsu, T. A Small Interfering RNA Targeting Vascular Endothelial Growth Factor as Cancer Therapeutics. Cancer Res. 64, 3365–3370 (2004).

21. Geisbert, T. W. et al. Postexposure Protection of Guinea Pigs against a Lethal Ebola Virus Challenge Is Conferred by RNA Interference. J. Infect. Dis. 193, 1650–1657 (2006).

22. Wolfrum, C. et al. Mechanisms and optimization of in vivo delivery of lipophilic siRNAs. Nat. Biotechnol. 25, 1149–1157 (2007).

23. Kumar, P. et al. Transvascular delivery of small interfering RNA to the central nervous system. Nature 448, 39–43 (2007).

24. Li, W. & Szoka, F. C. Lipid-based Nanoparticles for Nucleic Acid Delivery. Pharm. Res. 24, 438–449 (2007).

25. Morris, K. V. siRNA-mediated transcriptional gene silencing: the potential mechanism and a possible role in the histone code. Cell. Mol. Life Sci. CMLS 62, 3057–3066 (2005).

26. Park, W.-S., Hayafune, M., Miyano-Kurosaki, N. & Takaku, H. Specific HIV-1 env gene silencing by small interfering RNAs in human peripheral blood mononuclear cells. Gene Ther. 10, 2046–2050 (2003).

27. Alhoot, M. A., Wang, S. M. & Sekaran, S. D. Inhibition of Dengue Virus Entry and Multiplication into Monocytes Using RNA Interference. PLoS Negl. Trop. Dis. 5, e1410 (2011).

28. Ge, Q. et al. RNA interference of influenza virus production by directly targeting mRNA for degradation and indirectly inhibiting all viral RNA transcription. Proc. Natl. Acad. Sci. 100, 2718–2723 (2003).

29. Korf, M., Jarczak, D., Beger, C., Manns, M. P. & Krüger, M. Inhibition of hepatitis C virus translation and subgenomic replication by siRNAs directed against highly conserved HCV sequence and cellular HCV cofactors. J. Hepatol. 43, 225–234 (2005).

30. Giulietti, M. et al. To accelerate the Zika beat: Candidate design for RNA interference-based therapy. Virus Res. 255, 133–140 (2018).

31. Garber, K. Alnylam launches era of RNAi drugs. Nat. Biotechnol. 36, 777–778 (2018).

32. Hoy, S. M. Patisiran: First Global Approval. Drugs 78, 1625–1631 (2018).

33. Second RNAi drug approved. Nat. Biotechnol. 38, 385–385 (2020).

34. Scott, L. J. Givosiran: First Approval. Drugs 80, 335–339 (2020).

35. Mack, G. S. Erratum: MicroRNA gets down to business. Nat. Biotechnol. 29, 459–459 (2011).

36. Sayers, E. W. et al. Database resources of the National Center for Biotechnology Information. Nucleic Acids Res. 47, D23–D28 (2019).

37. Tamura, K. & Nei, M. Estimation of the number of nucleotide substitutions in the control region of mitochondrial DNA in humans and chimpanzees. Mol. Biol. Evol. 10, 512–526 (1993).

38. Kumar, S., Stecher, G. & Tamura, K. MEGA7: molecular evolutionary genetics analysis version 7.0 for bigger datasets. Mol. Biol. Evol. 33, 1870–1874 (2016).

39. Crooks, G. E., Hon, G., Chandonia, J.-M. & Brenner, S. E. WebLogo: a sequence logo generator. Genome Res. 14, 1188–1190 (2004).

40. Naito, Y., Yoshimura, J., Morishita, S. & Ui-Tei, K. siDirect 2.0: updated software for designing functional siRNA with reduced seed-dependent off-target effect. BMC Bioinformatics 10, 392 (2009).

41. Amarzguioui, M. & Prydz, H. An algorithm for selection of functional siRNA sequences. Biochem. Biophys. Res. Commun. 316, 1050–1058 (2004).

42. Lander, E. S. et al. Initial sequencing and analysis of the human genome. Nature 409, 860–921 (2001).

43. Ui-Tei, K., Naito, Y., Nishi, K., Juni, A. & Saigo, K. Thermodynamic stability and Watson– Crick base pairing in the seed duplex are major determinants of the efficiency of the siRNA-based off-target effect. Nucleic Acids Res. 36, 7100–7109 (2008).

44. Ui□Tei, K. et al. Guidelines for the selection of highly effective siRNA sequences for mammalian and chick RNA interference. Nucleic Acids Res. 32, 936–948 (2004).

45. Reynolds, A. et al. Rational siRNA design for RNA interference. Nat. Biotechnol. 22, 326–330 (2004).

46. Bernhart, S. H. et al. Partition function and base pairing probabilities of RNA heterodimers. Algorithms Mol. Biol. 1, 3 (2006).

47. Markham, N. R. & Zuker, M. DINAMelt web server for nucleic acid melting prediction. Nucleic Acids Res. 33, W577–W581 (2005).

48. Lu, Z. J., Gloor, J. W. & Mathews, D. H. Improved RNA secondary structure prediction by maximizing expected pair accuracy. RNA 15, 1805–1813 (2009).

49. Bellaousov, S., Reuter, J. S., Seetin, M. G. & Mathews, D. H. RNAstructure: web servers for RNA secondary structure prediction and analysis. Nucleic Acids Res. 41, W471–W474 (2013).

50. Rother, M. et al. ModeRNA server: an online tool for modeling RNA 3D structures. Bioinformatics 27, 2441–2442 (2011).

51. Krieger, E. et al. Improving physical realism, stereochemistry, and side-chain accuracy in homology modeling: Four approaches that performed well in CASP8. Proteins Struct. Funct. Bioinforma. 77, 114–122 (2009).

52. Cornell, W. D. et al. A Second Generation Force Field for the Simulation of Proteins, Nucleic Acids, and Organic Molecules J. Am. Chem. Soc. 1995, 117, 5179–5197. J. Am. Chem. Soc. 118, 2309–2309 (1996).

53. Yan, Y., Zhang, D., Zhou, P., Li, B. & Huang, S.-Y. HDOCK: a web server for protein– protein and protein–DNA/RNA docking based on a hybrid strategy. Nucleic Acids Res. 45, W365–W373 (2017).

54. Chan, C. Y. et al. A structural interpretation of the effect of GC-content on efficiency of RNA interference. BMC Bioinformatics 10, S33 (2009).

55. Aftab, S. O. et al. Analysis of SARS-CoV-2 RNA-dependent RNA polymerase as a potential therapeutic drug target using a computational approach. J. Transl. Med. 18, 275 (2020).

56. Ding, B., Qin, Y. & Chen, M. Nucleocapsid proteins: roles beyond viral RNA packaging. WIREs RNA 7, 213–226 (2016).

57. Schneider, T. D. & Stephens, R. M. Sequence logos: a new way to display consensus sequences. Nucleic Acids Res. 18, 6097–6100 (1990).

58. Chowdhury, U. F. et al. A computational approach to design potential siRNA molecules as a prospective tool for silencing nucleocapsid phosphoprotein and surface glycoprotein gene of SARS-CoV-2. Genomics 113, 331–343 (2021).

59. Sharif Shohan, M. U., Paul, A. & Hossain, M. Computational design of potential siRNA molecules for silencing nucleoprotein gene of rabies virus. Future Virol. 13, 159–170 (2018).

60. Mückstein, U. et al. Thermodynamics of RNA–RNA binding. Bioinformatics 22, 1177–1182 (2006).

61. Nur, S. M., Hasan, Md. A., Amin, M. A., Hossain, M. & Sharmin, T. Design of Potential RNAi (miRNA and siRNA) Molecules for Middle East Respiratory Syndrome Coronavirus (MERS-CoV) Gene Silencing by Computational Method. Interdiscip. Sci. Comput. Life Sci. 7, 257–265 (2015).

62. Kumar, M., Lata, S. & Raghava, G. siRNApred: SVM based method for predicting efficacy value of siRNA. in Proceedings of the OSCADD-2009 (2009).

63. Ding, Y., Chan, C. Y. & Lawrence, C. E. RNA secondary structure prediction by centroids in a Boltzmann weighted ensemble. RNA 11, 1157–1166 (2005).

64. Patzel, V. et al. Design of siRNAs producing unstructured guide-RNAs results in improved RNA interference efficiency. Nat. Biotechnol. 23, 1440–1444 (2005).

65. Neumeier, J. & Meister, G. siRNA Specificity: RNAi Mechanisms and Strategies to Reduce Off-Target Effects. Front. Plant Sci. 11, (2021).

66. Rand, T. A., Petersen, S., Du, F. & Wang, X. Argonaute2 Cleaves the Anti-Guide Strand of siRNA during RISC Activation. Cell 123, 621–629 (2005).

67. Krieger, E. & Vriend, G. New ways to boost molecular dynamics simulations. J. Comput. Chem. 36, 996–1007 (2015).

68. Land, H. & Humble, M. S. YASARA: A Tool to Obtain Structural Guidance in Biocatalytic Investigations. in Protein Engineering: Methods and Protocols (eds. Bornscheuer, U. T. & Höhne, M.) 43–67 (Springer, 2018). doi:10.1007/978-1-4939-7366-8_4.

69. Thompson, E., Thackray, T., Kalthoff, C., Rapoport, R. & Jagodzinski, F. Using Energy-Minimization Profiles to Measure Protein Resistance to Drugs. in Proceedings of the 11th ACM International Conference on Bioinformatics, Computational Biology and Health Informatics 1–6 (Association for Computing Machinery, 2020). doi:10.1145/3388440.3414703.

70. Gautam, B. Energy Minimization. Homol. Mol. Model. -Perspect. Appl. (2020) doi:10.5772/intechopen.94809.

71. Roy, K., Kar, S. & Das, R. N. Chapter 5 -Computational Chemistry. in Understanding the Basics of QSAR for Applications in Pharmaceutical Sciences and Risk Assessment (eds. Roy, K., Kar, S. & Das, R. N.) 151–189 (Academic Press, 2015). doi:10.1016/B978-0-12-801505-6.00005-3.

72. Mackay, D. H. J., Cross, A. J. & Hagler, A. T. The Role of Energy Minimization in Simulation Strategies of Biomolecular Systems. in Prediction of Protein Structure and the Principles of Protein Conformation (ed. Fasman, G. D.) 317–358 (Springer US, 1989). doi:10.1007/978-1-4613-1571-1_7.

73. Hutvagner, G. & Simard, M. J. Argonaute proteins: key players in RNA silencing. Nat. Rev. Mol. Cell Biol. 9, 22–32 (2008).

74. Song, Y. et al. COVID-19 treatment: close to a cure? A rapid review of pharmacotherapies for the novel coronavirus (SARS-CoV-2). Int. J. Antimicrob. Agents 56, 106080 (2020).

75. Takahashi, T. et al. Interactions between the non-seed region of siRNA and RNA-binding RLC/RISC proteins, Ago and TRBP, in mammalian cells. Nucleic Acids Res. 42, 5256–5269 (2014).

76. Bhandare, V. & Ramaswamy, A. Structural Dynamics of Human Argonaute2 and Its Interaction with siRNAs Designed to Target Mutant tdp43. Adv. Bioinforma. 2016, e8792814 (2016).

77. Elkayam, E. et al. The Structure of Human Argonaute-2 in Complex with miR-20a. Cell 150, 100–110 (2012).

78. Hasan, M. et al. Computational prediction of potential siRNA and human miRNA sequences to silence orf1ab associated genes for future therapeutics against SARS-CoV-2. Inform. Med. Unlocked 24, 100569 (2021).

79. Chen, Y., Cheng, G. & Mahato, R. I. RNAi for Treating Hepatitis B Viral Infection. Pharm. Res. 25, 72–86 (2008).

