## Supplementary for "SiRNA Molecules as Potential RNAi Therapeutics to Silence RdRP Region and N-Gene of SARS-CoV-2: An *In Silico* Approach": Supplementary.docx

**Supplementary Data**

**Table S1.** **Algorithms or rules for the rational design of siRNA molecules**

| **E-value**  **E = Kmn e^-λS^**  K, **λ =** high-scoring segment pairs parameters  m, n = limit of sufficiently large sequence lengths | | |
| --- | --- | --- |
| **Ui-Tei rules** | **Amarzguioui rules** | **Reynolds rules** |
| - A/U at the 5′ terminus of the sense strand - G/C at the 5′ terminus of the antisense strand - At least 4 A/U residues in the 5′ terminal 7 bp of sense strand - No GC stretch longer than 9 nucleotides | - Duplex End A/U differential >0 - Strong binding of 5′ sense strand - No U at position 1 - Presence of A at position 6 - Weak binding of 3′ sense strand | - GC content 30–52% (one point) - Occurrence of three or more A/U base pair at position 15–19 of the sense strand (Each A/U base pair in this region earns one point) - Low internal stability at the target site (Tm>−20◦C) (one point) - Presence of A at position 19 of the sense strand (one point) - Presence of A at position three of the sense strand (one point) - Presence of U at position ten of the sense strand (one point) - Absence of G at position 13 of the sense strand (one point) - Threshold for efficient siRNAs score >= 6 |
| T_m_ calculation formula  T_m_ = {(1000× ∆H)/ (A+∆S+R ln (CT/4)) - 273.15 + 16.6 log [Na+]  Here,  ∆H = the sum of the nearest neighbor enthalpy change (kcal/ mol).  A = the helix initiation constant (-10.8).  ∆S = the sum of the nearest neighbor entropy change.  R = the gas constant (1.987 cal/deg/mol).  CT = the total molecular concentration of the strand (100 μM).  [Na+] was fixed at 100 mM. | | |
| Free energy of the heterodimer of sequence A and sequence B  ∆Gbinding = ∆GAB - ∆GA - ∆GB | | |

**Table S2. Conserved sequences and their corresponding positions for RdRP region of different SARS-CoV-2 variants**

| **Positions** | **Sequences** |
| --- | --- |
| 1-94 | TCAGCTGATGCACAATCGTTTTTAAACGGGTTTGCGGTGTAAGTGCAGCCCGTCTTACACCGTGCGGCACAGGCACTAGTACTGATGTCGTATA |
| 96-178 | AGGGCTTTTGACATCTACAATGATAAAGTAGCTGGTTTTGCTAAATTCCTAAAAACTAATTGTTGTCGCTTCCAAGAAAAGGA |
| 180-205 | GAAGATGACAATTTAATTGATTCTTA |
| 207-418 | TTTGTAGTTAAGAGACACACTTTCTCTAACTACCAACATGAAGAAACAATTTATAATTTACTTAAGGATTGTCCAGCTGTTGCTAAACATGACTTCTTTAAGTTTAGAATAGACGGTGACATGGTACCACATATATCACGTCAACGTCTTACTAAATACACAATGGCAGACCTCGTCTATGCTTTAAGGCATTTTGATGAAGGTAATTGTGA |
| 420-550 | ACATTAAAAGAAATACTTGTCACATACAATTGTTGTGATGATGATTATTTCAATAAAAAGGACTGGTATGATTTTGTAGAAAACCCAGATATATTACGCGTATACGCCAACTTAGGTGAACGTGTACGCCA |
| 552-678 | GCTTTGTTAAAAACAGTACAATTCTGTGATGCCATGCGAAATGCTGGTATTGTTGGTGTACTGACATTAGATAATCAAGATCTCAATGGTAACTGGTATGATTTCGGTGATTTCATACAAACCACGC |
| 680-745 | AGGTAGTGGAGTTCCTGTTGTAGATTCTTATTATTCATTGTTAATGCCTATATTAACCTTGACCAG |
| 747-820 | GCTTTAACTGCAGAGTCACATGTTGACACTGACTTAACAAAGCCTTACATTAAGTGGGATTTGTTAAAATATGA |
| 822-920 | TTCACGGAAGAGAGGTTAAAACTCTTTGACCGTTATTTTAAATATTGGGATCAGACATACCACCCAAATTGTGTTAACTGTTTGGATGACAGATGCATT |
| 922-966 | TGCATTGTGCAAACTTTAATGTTTTATTCTCTACAGTGTTCCCAC |
| 968-1234 | TACAAGTTTTGGACCACTAGTGAGAAAAATATTTGTTGATGGTGTTCCATTTGTAGTTTCAACTGGATACCACTTCAGAGAGCTAGGTGTTGTACATAATCAGGATGTAAACTTACATAGCTCTAGACTTAGTTTTAAGGAATTACTTGTGTATGCTGCTGACCCTGCTATGCACGCTGCTTCTGGTAATCTATTACTAGATAAACGCACTACGTGCTTTTCAGTAGCTGCACTTACTAACAATGTTGCTTTTCAAACTGTCAAACC |
| 1236-1344 | GGTAATTTTAACAAAGACTTCTATGACTTTGCTGTGTCTAAGGGTTTCTTTAAGGAAGGAAGTTCTGTTGAATTAAAACACTTCTTCTTTGCTCAGGATGGTAATGCTG |
| 1365-1815 | TATCGTTATAATCTACCAACAATGTGTGATATCAGACAACTACTATTTGTAGTTGAAGTTGTTGATAAGTACTTTGATTGTTACGATGGTGGCTGTATTAATGCTAACCAAGTCATCGTCAACAACCTAGACAAATCAGCTGGTTTTCCATTTAATAAATGGGGTAAGGCTAGACTTTATTATGATTCAATGAGTTATGAGGATCAAGATGCACTTTTCGCATATACAAAACGTAATGTCATCCCTACTATAACTCAAATGAATCTTAAGTATGCCATTAGTGCAAAGAATAGAGCTCGCACCGTAGCTGGTGTCTCTATCTGTAGTACTATGACCAATAGACAGTTTCATCAAAAATTATTGAAATCAATAGCCGCCACTAGAGGAGCTACTGTAGTAATTGGAACAAGCAAATTCTATGGTGGTTGGCACAACATGTTAAAAACTGTTT |
| 1839-1882 | CTTATGGGTTGGGATTATCCTAAATGTGATAGAGCCATGCCTAA |
| 1884-2009 | ATGCTTAGAATTATGGCCTCACTTGTTCTTGCTCGCAAACATACAACGTGTTGTAGCTTGTCACACCGTTTCTATAGATTAGCTAATGAGTGTGCTCAAGTATTGAGTGAAATGGTCATGTGTGGC |
| 2011-2465 | GTTCACTATATGTTAAACCAGGTGGAACCTCATCAGGAGATGCCACAACTGCTTATGCTAATAGTGTTTTTAACATTTGTCAAGCTGTCACGGCCAATGTTAATGCACTTTTATCTACTGATGGTAACAAAATTGCCGATAAGTATGTCCGCAATTTACAACACAGACTTTATGAGTGTCTCTATAGAAATAGAGATGTTGACACAGACTTTGTGAATGAGTTTTACGCATATTTGCGTAAACATTTCTCAATGATGATACTCTCTGACGATGCTGTTGTGTGTTTCAATAGCACTTATGCATCTCAAGGTCTAGTGGCTAGCATAAAGAACTTTAAGTCAGTTCTTTATTATCAAAACAATGTTTTTATGTCTGAAGCAAAATGTTGGACTGAGACTGACCTTACTAAAGGACCTCATGAATTTTGCTCTCAACATACAATGCTAGTTAAACAG |
| 2485-2530 | TTCCTTACCCAGATCCATCAAGAATCCTAGGGGCCGGCTGTTTTGT |
| 2532-2602 | GATGATATCGTAAAAACAGATGGTACACTTATGATTGAACGGTTCGTGTCTTTAGCTATAGATGCTTACCC |
| 2604-2734 | CTTACTAAACATCCTAATCAGGAGTATGCTGATGTCTTTCATTTGTACTTACAATACATAAGAAAGCTACATGATGAGTTAACAGGACACATGTTAGACATGTATTCTGTTATGCTTACTAATGATAACAC |
| 2736-2795 | CTTACTAAACATCCTAATCAGGAGTATGCTGATGTCTTTCATTTGTACTTACAATACATAAGAAAGCTACATGATGAGTTAACAGGACACATGTTAGACATGTATTCTGTTATGCTTACTAATGATAACAC |

**Table S3. Conserved sequences and their corresponding positions for N-gene region of different SARS-CoV-2 variants**

| **Positions** | **Sequences** |
| --- | --- |
| 10-122 | AATGGACCCCAAAATCAGCGAAATGCACCCCGCATTACGTTTGG TGGACCCTCAGATTCAACTGGCAGTAACCAGAATGGAGAACGCA  GTGGGGCGCGATCAAAACAACGTCG |
| 124-187 | CCCCAAGGTTTACCCAATAATACTGCGTCTTGGTTCACCGCTCTCA  CTCAACATGGCAAGGAAG |
| 189-238 | CCTTAAATTCCCTCGAGGACAAGGCGTTCCAATTAACACCAATAG  CAGTC |
| 240-368 | AGATGACCAAATTGGCTACTACCGAAGAGCTACCAGACGAATTC  GTGGTGGTGACGGTAAAATGAAAGATCTCAGTCCAAGATGGTAT  TTCTACTACCTAGGAACTGGGCCAGAAGCTGGACTTCCCTA |
| 418-485 | AATACACCAAAAGATCACATTGGCACCCGCAATCCTGCTAACAA  TGCTGCAATCGTGCTACAACTTCC |
| 487-518 | CAAGGAACAACATTGCCAAAAGGCTTCTACGC |
| 520-580 | GAAGGGAGCAGAGGCGGCAGTCAAGCCTCTTCTCGTTCCTCATC  ACGTAGTCGCAACAGTT |
| 615-703 | TTCTCCTGCTAGAATGGCTGGCAATGGCGGTGATGCTGCTCTTGC  TTTGCTGCTGCTTGACAGATTGAACCAGCTTGAGAGCAAAATGT |
| 705-806 | TGGTAAAGGCCAACAACAACAAGGCCAAACTGTCACTAAGAAAT  CTGCTGCTGAGGCTTCTAAGAAGCCTCGGCAAAAACGTACTGCC  ACTAAAGCATACAA |
| 808-870 | GTAACACAAGCTTTCGGCAGACGTGGTCCAGAACAAACCCAAGG  AAATTTTGGGGACCAGGAA |
| 876-998 | CAGACAAGGAACTGATTACAAACATTGGCCGCAAATTGCACAAT  TTGCCCCCAGCGCTTCAGCGTTCTTCGGAATGTCGCGCATTGGCA  TGGAAGTCACACCTTCGGGAACGTGGTTGACCTA |
| 1000-1084 | ACAGGTGCCATCAAATTGGATGACAAAGATCCAAATTTCAAAGA  TCAAGTCATTTTGCTGAATAAGCATATTGACGCATACAAAA |
| 1110-1177 | GGACAAAAAGAAGAAGGCTGATGAAACTCAAGCCTTACCGCAGA  GACAGAAGAAACAGCAAACTGTGA |
| 1194-1252 | AGATTTGGATGATTTCTCCAAACAATTGCAACAATCCATGAGCAG  TGCTGACTCAACTC |

**Table S4. Effective siRNA molecules against RdRP region with T_m_ values of both guide strand and passenger strand, guide strands’ GC%, free energy of binding (MEF) with target, T_m_C_p,_ T_m_(Conc), and validity (binary).**

| **Location of target within mRNA** | **siRNA target within mRNA** | **Predicted siRNA duplex candidates at 37^o^ C**  **(21nt guide (5′→3′)**  **21nt passenger (5′→3′))** | **Seed-duplex stability (T_m_)^o^C** | | **GC% (%)** | **MFE**  **(Kcal/mol)** | **T_m_C_p_**  **°C** | **T_m_(conc)**  **°C** | **Validity (binary)** |
| --- | --- | --- | --- | --- | --- | --- | --- | --- | --- |
|  |  |  | **Guide** | **Passenger** |  |  |  |  |  |
| 228-250 | TTCTCTAACTACCAACATGAAGA | UUCAUGUUGGUAGUUAGAGAA  CUCUAACUACCAACAUGAAGA | 20.5°C | 18.9°C | 33.33 | -32.7 | 81.2 | 79.9 | 1.018 |
| 585-607 | ATGCGAAATGCTGGTATTGTTGG | AACAAUACCAGCAUUUCGCAU  GCGAAAUGCUGGUAUUGUUGG | 14.6°C | 14.1°C | 38.1 | -33.6 | 84.3 | 83.2 | 1.019 |
| 1267-1289 | CTGTGTCTAAGGGTTTCTTTAAG | UAAAGAAACCCUUAGACACAG  GUGUCUAAGGGUUUCUUUAAG | 5.5°C | 20.3°C | 38.1 | -33.9 | 83.8 | 82.4 | 0.959 |
| 1378-1400 | TACCAACAATGTGTGATATCAGA | UGAUAUCACACAUUGUUGGUA  CCAACAAUGUGUGAUAUCAGA | 17.4°C | 12.1°C | 33.33 | -32.9 | 79.9 | 79.0 | 0.963 |
| 1573-1595 | ATGCACTTTTCGCATATACAAAA | UUGUAUAUGCGAAAAGUGCAU  GCACUUUUCGCAUAUACAAAA | 8.2°C | 10.3°C | 33.33 | -31.7 | 80.5 | 81.5 | 1.003 |
| 1636-1658 | ATGCCATTAGTGCAAAGAATAGA | UAUUCUUUGCACUAAUGGCAU  GCCAUUAGUGCAAAGAAUAGA | 5.3°C | 17.4°C | 33.33 | -32.5 | 83.0 | 81.7 | 0.970 |
| 1675-1697 | GTGTCTCTATCTGTAGTACTATG | UAGUACUACAGAUAGAGACAC  GUCUCUAUCUGUAGUACUAUG | 18.8°C | 20.2°C | 38.1 | -35.2 | 82.8 | 81.5 | 1.007 |
| 1688-1710 | TAGTACTATGACCAATAGACAGT | UGUCUAUUGGUCAUAGUACUA  GUACUAUGACCAAUAGACAGU | 14.5°C | 13.1°C | 33.33 | -33.5 | 81.3 | 80.3 | 0.969 |
| 1697-1719 | GACCAATAGACAGTTTCATCAAA | UGAUGAAACUGUCUAUUGGUC  CCAAUAGACAGUUUCAUCAAA | 16.3°C | 13.4°C | 38.1 | -34.5 | 80.6 | 79.4 | 0.960 |
| 2414-2436 | TACTAAAGGACCTCATGAATTTT | AAUUCAUGAGGUCCUUUAGUA  CUAAAGGACCUCAUGAAUUUU | 13.6°C | 19.9°C | 33.33 | -32.1 | 82.8 | 81.5 | 0.985 |
| 2622-2644 | CAGGAGTATGCTGATGTCTTTCA | AAAGACAUCAGCAUACUCCUG  GGAGUAUGCUGAUGUCUUUCA | 19.2°C | 20.3°C | 42.86 | -35.9 | 85.1 | 83.7 | 0.976 |
| 2667-2689 | AAGCTACATGATGAGTTAACAGG | UGUUAACUCAUCAUGUAGCUU  GCUACAUGAUGAGUUAACAGG | 12.9°C | 17.9°C | 33.33 | -32.7 | 81.1 | 79.9 | 1.015 |
| 2679-2701 | GAGTTAACAGGACACATGTTAGA | UAACAUGUGUCCUGUUAACUC  GUUAACAGGACACAUGUUAGA | 19.3°C | 11.8°C | 38.1 | -34.3 | 82.2 | 81.1 | 0.951 |

**Table S5. Effective siRNA molecules against N-gene with T_m_ values of both guide strand and passenger strand, guide strands’ GC%, free energy of binding (MEF) with target, T_m_C_p,_ T_m_(Conc), and validity (binary).**

| **Location of target within mRNA** | **siRNA target within mRNA** | **Predicted siRNA duplex candidates at 37^o^ C**  **21nt guide (5′→3′)**  **21nt passenger (5′→3′)** | **Seed-duplex stability (T_m_)°C** | | **GC% (%)** | **MFE**  **(Kcal/mol)** | **T_m_C_p_**  **°C** | **T_m_(conc)**  **°C** | **Validity (binary)** |
| --- | --- | --- | --- | --- | --- | --- | --- | --- | --- |
|  |  |  | **Guide** | **Passenger** |  |  |  |  |  |
| 314-336 | GTCCAAGATGGTATTTCTACTAC | AGUAGAAAUACCAUCUUGGAC  CCAAGAUGGUAUUUCUACUAC | 14.6°C | 18.1°C | 38.1 | -34.5 | 80.7 | 80.1 | 0.985 |
| 411-433 | AGCCTTGAATACACCAAAAGATC | UCUUUUGGUGUAUUCAAGGCU  CCUUGAAUACACCAAAAGAUC | 16.1°C | 12.0°C | 38.1 | -34.2 | 82.4 | 81.2 | 0.986 |
| 727-749 | GGCCAAACTGTCACTAAGAAATC | UUUCUUAGUGACAGUUUGGCC  CCAAACUGUCACUAAGAAAUC | 11.7°C | 16.7°C | 42.86 | -35.8 | 83.9 | 82.6 | 0.959 |
| 789-811 | TGCCACTAAAGCATACAATGTAA | ACAUUGUAUGCUUUAGUGGCA  CCACUAAAGCAUACAAUGUAA | 13.5°C | 11.8°C | 38.1 | -34.5 | 82.9 | 82.1 | 0.986 |
| 836-858 | CAGAACAAACCCAAGGAAATTTT | AAUUUCCUUGGGUUUGUUCUG  GAACAAACCCAAGGAAAUUUU | 18.7°C | 13.3°C | 38.1 | -32.5 | 82.7 | 81.6 | 0.934 |
| 892-914 | TACAAACATTGGCCGCAAATTGC | AAUUUGCGGCCAAUGUUUGUA  CAAACAUUGGCCGCAAAUUGC | 20.6°C | 5.3°C | 38.1 | -33.1 | 85.5 | 84.4 | 0.950 |
| 1063-1085 | AAGCATATTGACGCATACAAAAC | UUUGUAUGCGUCAAUAUGCUU  GCAUAUUGACGCAUACAAAAC | 13.5°C | 5.6°C | 33.33 | -31.7 | 81.8 | 80.9 | 1.047 |

**Table S6. Docking interaction between receptor-ligand (Ago2-siRNA) interface residue pairs for selected siRNAs against the RdRP and N regions of SARS-CoV-2. [rN, N = serial number of RdRP targeting siRNAs and nN, N = serial number of N-gene targeting siRNAs]**

| **Alias** | **Receptor-ligand interface residue pair(s)** |
| --- | --- |
| **r1** | LYS65-17A, LYS65-18A, CYS66-17A, PRO67-16G, PRO67-17A, ARG68-14G, ARG68-15A, ARG68-16G, ARG69-15A, VAL70-17A, ASP95-14G, ARG97-14G, ARG97-15A, ARG97-16G, PRO120-18A, PHE128-17A, PHE156-11A, PHE156-12U, GLN160-12U, ARG167-13A, PRO176-14G, PRO176-15A, VAL177-14G, GLY178-13A, GLY178-14G, ARG179-11A, ARG179-13A, ARG179-14G, VAL219-8U, SER220-8U, ALA221-7U, ALA221-8U, THR222-8U, THR222-9G, ALA223-8U, LYS266-15A, LYS266-16G, LYS266-17A, ARG277-17A, ARG277-18A, LYS278-17A, LYS278-18A, TYR279-17A, TYR279-18A, ARG280-16G, ARG280-17A, ARG351-8U, ARG351-9G, ARG351-10G, ARG351-14G, CYS352-9G, ILE353-9G, ILE353-10G, ILE353-14G, LYS354-9G, LYS355-9G, LYS355-10G, LEU356-8U, LEU356-9G, THR361-6G, THR361-7U, THR361-8U, SER362-6G, MET364-7U, MET364-8U, ILE365-6G, ILE365-7U, ARG366-6G, THR368-7U, ALA369-6G, ALA369-7U, ARG375-7U, GLY433-5U, LEU522-1U, GLY524-1U, LYS525-1U, THR526-1U, TYR529-1U, LYS533-1U, THR544-1U, GLN545-1U, GLN545-2U, CYS546-1U, CYS546-2U, VAL547-1U, VAL547-2U, GLN548-1U, GLN548-2U, ASN551-1U, ASN551-2U, GLN558-2U, THR559-2U, ASN562-2U, ASN562-3C, LEU563-2U, LEU563-3C, LYS566-1U, LYS566-2U, LYS566-3C, LYS570-1U, ASP597-10G, VAL598-10G, VAL598-11U, THR599-10G, HIS600-10G, HIS600-11U, ARG635-10G, ARG635-11U, ARG635-11A, ARG635-13A, SER672-11U, SER672-11A, GLU673-11A, GLU673-11G, GLU673-11U, GLY674-11A, GLY674-11G, GLY674-11U, GLY674-12U, GLN675-11U, GLN675-11A, GLN675-11G, GLN675-11U, PHE676-11G, GLN677-11G, GLN678-11G, LYS709-6G, ARG710-8U, ARG710-9G, ARG714-6G, ARG714-7U, ARG714-8U, HIS753-4A, HIS753-5U, HIS753-6G, ALA754-4A, ALA754-5U, GLY755-5U, ILE756-4A, ILE756-5U, GLN757-5U, GLN757-6G, GLY758-6G, THR759-6G, THR759-7U, SER760-5U, SER760-6G, ARG761-5U, ARG761-6G, ARG761-7U, PRO762-6G, TYR790-3C, TYR790-4A, ARG792-1U, ARG792-2U, ARG792-3C, ARG792-4A, CYS793-3C, CYS793-4A, THR794-4A, ARG795-3C, ARG795-4A, ARG795-5U, SER796-4A, SER796-5U, VAL797-4A, VAL797-5U, SER798-4A, SER798-5U, SER798-6G, HIS807-10G, ARG812-1U, TYR815-1U, ALA859-1U |
| **r2** | SER362-11C, ILE365-11C, ARG366-11C, ARG366-11A, ARG366-11G, SER371-12C, ASP374-12C, GLU378-12C, LEU522-3A, GLY524-3A, LYS525-3A, THR526-3A, TYR529-3A, GLN545-3A, GLN545-4A, CYS546-3A, CYS546-4A, VAL547-3A, GLN548-3A, GLN558-6A, GLN558-7A, THR559-6A, THR559-7A, ASN562-5C, ASN562-6A, LEU563-4A, LYS566-3A, LYS566-4A, LYS566-5C, LYS720-14U, LYS720-14U, ASN721-14U, ARG723-14U, ARG723-14U, GLY725-14U, LYS726-10C, LYS726-11C, LYS726-12C, LYS726-13A, ILE756-6A, ILE756-7A, ILE756-8U, TYR790-5C, TYR790-6A, ARG792-4A, ARG792-5C, ARG792-6A, CYS793-5C, CYS793-6A, ARG795-6A, ARG795-7A, VAL797-6A, TYR804-5C, TYR804-6A, LEU808-4A, LEU808-5C, PHE811-5C, ARG812-3A, TYR815-3A, TYR815-4A, ALA859-4A |
| **r3** | LYS65-17C, LYS65-17A, CYS66-17C, PRO67-16A, PRO67-17C, ARG68-14A, ARG68-15G, ARG69-15G, ARG97-14A, ARG97-15G, GLY178-13U, GLY178-14A, ARG179-12U, ARG179-13U, ALA221-8A, ARG277-17G, TYR279-17A, TYR279-17G, ARG315-17G, LYS335-17A, ARG351-8A, ARG351-9C, THR361-7A, MET364-7A, ILE365-6A, ILE365-7A, ARG375-6A, ARG375-7A, LEU522-1U, GLY524-1U, LYS525-1U, THR526-1U, TYR529-1U, ALA530-1U, LYS533-1U, THR544-1U, GLN545-1U, CYS546-1U, CYS546-2A, VAL547-1U, VAL547-2A, GLN548-1U, GLN548-2A, ASN551-1U, ASN551-2A, GLN558-2A, THR559-2A, ASN562-2A, ASN562-3A, LEU563-2A, LYS566-1U, LYS566-2A, LYS566-3A, LYS570-1U, ASP597-10C, VAL598-10C, THR599-10C, HIS600-10C, HIS600-11C, PRO601-9C, PRO601-10C, PRO601-11C, PRO602-9C, PRO602-10C, ALA603-9C, ALA603-10C, GLY604-10C, ARG635-10C, ARG635-11C, ARG635-12U, GLU637-11C, GLY670-11C, VAL671-11C, SER672-11C, SER672-12U, GLY674-12U, GLN675-11C, GLN675-12U, GLN678-12U, LYS709-5G, ARG710-8A, ARG710-9C, ARG710-10C, ARG714-7A, ASN729-6A, HIS753-5G, HIS753-6A, ALA754-4A, ALA754-5G, GLY755-5G, GLY755-6A, ILE756-4A, ILE756-5G, GLN757-5G, GLN757-6A, GLY758-5G, GLY758-6A, THR759-6A, SER760-5G, SER760-6A, ARG761-6A, ARG761-7A, ARG761-8A, TYR790-3A, TYR790-4A, ARG792-1U, ARG792-3A, ARG792-4A, CYS793-3A, CYS793-4A, ARG795-3A, ARG795-4A, VAL797-4A, SER798-5G, TYR804-4A, TYR804-5G, HIS807-10C, PHE811-10C, ARG812-1U, TYR815-1U, ALA859-1U, ALA859-3A |
| **r4** | LYS355-0A, LYS355-0A, ASP358-1U, ASN359-9G, SER362-9G, THR363-9G, ARG366-9G, ARG366-10U, GLN548-3C, GLN548-4A, ASN551-2U, ASN551-3C, ASN551-4A, ARG554-4A, ARG554-5U, THR556-3C, THR556-4A, GLN558-3C, GLN558-4A, THR559-2U, THR559-3C, THR559-4A, ASN562-2U, ASN562-3C, ASN562-4A, LEU563-2U, LYS566-2U, LYS709-0A, LYS709-0A, ARG710-0A, ARG761-0A, ARG761-0A, TYR790-2U, ARG792-1U, ARG792-2U, ARG792-3C, CYS793-2U, CYS793-3C, ARG795-4A, TYR804-0A, TYR804-1U, HIS807-0A, HIS807-0A, LEU808-1U, LEU808-2U, PHE811-0A, PHE811-1U |
| **r5** | GLU58-8C, GLU58-9A, LYS65-15C, PRO67-15C, PRO67-16U, ARG68-14A, ARG68-15C, ARG68-16U, ARG69-16U, ARG69-17C, ASP95-9A, ARG97-9A, ARG97-14A, LYS98-8C, LYS98-9A, ASN99-8C, ASN99-9A, GLN141-0A, ALA142-0A, ALA142-1A, ASP145-0A, ALA146-0A, ARG150-0A, LEU151-0A, LEU151-1A, PRO152-1A, PHE156-10G, GLN160-10G, TYR174-17C, THR175-16U, THR175-17C, PRO176-15C, PRO176-16U, PRO176-17C, VAL177-16U, ALA184-17C, TRP199-17C, TRP199-17C, PHE224-16U, ALA227-17C, LYS266-15C, LYS266-16U, GLU268-17C, LYS278-17C, LYS278-17C, ARG280-14A, ARG280-15C, VAL347-17C, ALA348-16U, ALA348-17C, ALA348-17C, GLY349-16U, GLY349-17C, GLN350-16U, ARG351-16U, ILE353-15C, ILE353-16U, HIS634-13U, ARG635-12A, ARG635-13U, SER672-11C, GLY674-10G, GLY674-11C, GLN675-11C, GLN675-12A, GLN678-10G |
| **n1** | TYR101-18C, PHE156-17A, PHE156-18C, GLU157-17A, GLU157-18C, GLN160-17A, GLN160-18C, ARG179-10C, ARG179-15G, ARG351-11C, CYS352-11C, ILE353-11C, ILE353-11A, ASP358-1A, SER362-1A, ILE365-1A, ARG366-1A, ALA367-1A, THR368-1A, ARG370-1A, LEU522-5A, PRO523-5A, GLY524-5A, LYS525-5G, LYS525-5A, TYR529-5A, LYS533-5A, CYS546-5A, VAL547-5A, GLN548-5A, LYS550-4A, ASP597-8U, HIS600-13U, PRO601-12C, PRO602-12C, ALA603-12C, GLY604-12C, ARG635-12C, ARG635-13U, ARG635-14U, GLU637-14U, GLU673-16G, GLY674-15G, GLY674-16G, GLN675-14U, GLN675-15G, GLN675-16G, GLN677-17A, GLN678-15G, LYS709-8U, ARG710-8U, ARG710-9A, ARG710-10C, ARG761-8U, ARG761-9A, TYR790-6A, ARG792-6A, TYR804-6A, TYR804-7A, HIS807-7A, HIS807-8U, LEU808-6A, LEU808-7A, PHE811-5A, PHE811-7A, TYR815-5A |
| **n2** | PRO67-15G, PRO67-16G, ARG68-13U, ARG68-14U, ARG68-15G, ASP95-13U, ASP95-13U, ASP95-14U, ARG97-13U, ARG97-14U, ARG97-15G, ASN99-13U, PHE156-13U, GLU157-13U, GLN160-13U, GLY178-13U, ARG179-12A, PHE182-8U, ASP218-7G, VAL219-7G, SER220-7G, SER220-8U, ALA221-7G. ALA221-8U, THR222-7G, THR222-8U, THR222-9G, ARG280-15G, ARG280-16G, ARG280-17C, GLN332-17C, GLN332-18C, LYS335-18C, ARG351-8U, ARG351-9G, ARG351-10A, ILE353-9G, THR361-7G, MET364-7G, ILE365-6A, ILE365-7G, THR368-6A, THR368-7G, ARG375-6A, ARG375-7G, LEU522-0U, GLY524-0U, GLY524-1U, LYS525-0U, LYS525-1U, TYR529-0U, LYS533-0U, GLN545-0U, GLN545-1U, CYS546-0U, CYS546-1U, VAL547-0U, VAL547-1U, GLN548-0U, GLN548-1U, LYS550-1U, ASN551-0U, ASN551-1U, ASN551-2U, GLN558-2U, THR559-2U, ASN562-2U, LEU563-1U, LYS566-0U, LYS566-1U, LYS570-0U, VAL598-10A, THR599-10A, HIS600-10A, HIS600-11C, PRO601-10A, PRO601-11C, PRO602-9G, PRO602-10A, PRO602-11C, ALA603-9G, ALA603-10A, GLY604-10A, HIS634-11C, ARG635-10A, ARG635-11C, ARG635-12A, GLN636-11C, GLU637-11C, GLY670-11C, SER672-11C, SER672-12A, GLN675-11C, GLN675-12A, LYS709-4U, LYS709-5U, LYS709-6A, ARG710-8U, ARG710-9G, ARG710-10A, ARG714-6A, ARG714-7G, ASN729-5U, HIS753-4U, HIS753-5U, ALA754-4U, ALA754-5U, GLY755-4U, GLY755-5U, ILE756-3C, ILE756-4U, ILE756-5U, GLN757-4U, GLN757-5U, GLN757-6A, GLY758-5U, GLY758-6A, THR759-5U, THR759-6A, SER760-5U, SER760-6A, ARG761-5U, ARG761-6A, ARG761-7G, PRO762-6A, TYR790-4U, ARG792-3C, CYS793-3C, CYS793-4U, ARG795-3C, ARG795-4U, VAL797-4U, SER798-4U, SER798-5U, TYR804-4U, PHE811-10A, ARG812-0U, TYR815-0U, PHE858-0U, ALA859-0U |
| **n3** | LYS65-17U, LYS65-17U, CYS66-17U, PRO67-16U, PRO67-17U, ARG68-14U, ARG68-15G, ARG68-16U, ARG97-13A, ARG97-14U, ARG97-15G, GLY178-13A, GLY178-14U, ARG179-12A, ARG179-13A, SER220-8G, ALA221-8G, ARG277-17A, TYR279-17U, TYR279-17A, ARG315-17A, GLN332-17U, GLN332-17G, ARG351-9G, ILE365-6G, ILE365-7C, ARG375-7C, LEU522-1A, GLY524-1A, LYS525-1A, THR526-1A, TYR529-1A, LYS533-1A, GLN545-1A, GLN545-2A, CYS546-1A, CYS546-2A, VAL547-1A, VAL547-2A, GLN548-1A, GLN548-2A, ASN551-1A, ASN551-2A, GLN558-2A, THR559-2A, ASN562-2A, ASN562-3U, LEU563-2A, LYS566-1A, LYS566-2A, LYS566-3U, LYS570-1A, VAL598-10C, THR599-10C, HIS600-10C, HIS600-11C, PRO601-9G, PRO601-10C, PRO601-11C, PRO602-9G, PRO602-10C, ALA603-9G, ALA603-10C, ARG635-10C, ARG635-11C, GLU637-11C, GLY670-11C, VAL671-11C, SER672-11C, SER672-12A, GLY674-12A, GLN675-11C, GLN675-12A, ARG710-9G, ARG710-10C, ARG710-11C, ARG710-12A, ASN729-5U, ASN729-6G, HIS753-4U, HIS753-5U, HIS753-6G, ALA754-4U, ALA754-5U, GLY755-5U, GLY755-6G, ILE756-4U, ILE756-5U, ILE756-6G, GLN757-5U, GLN757-6G, GLY758-5U, GLY758-6G, THR759-6G, SER760-5U, SER760-6G, ARG761-6G, ARG761-7C, ARG761-8G, TYR790-3U, TYR790-4U, ARG792-1A, ARG792-2A, ARG792-3U, ARG792-4U, CYS793-3U, CYS793-4U, ARG795-4U, ARG795-5U, SER796-5U, VAL797-4U, VAL797-5U, SER798-4U, SER798-5U, TYR804-3U, TYR804-4U, PHE811-9G, PHE811-10C, ARG812-1A, TYR815-1A, PHE858-1A, ALA859-1A, ALA859-3U |
